# Genetic and functional evidence relates a missense variant in *B4GALT1* to lower LDL-C and fibrinogen

**DOI:** 10.1101/721704

**Authors:** May E. Montasser, Cristopher V.Van Hout, Rebecca McFarland, Avraham Rosenberg, Myrasol Callaway, Biao Shen, Ning Li, Thomas J. Daly, Alicia D. Howard, Wei Lin, Yuan Mao, Bin Ye, Giusy Della Gatta, Gannie Tzoneva, James Perry, Kathleen A. Ryan, Lawrence Miloscio, Aris N. Economides, Regeneron Genetics Center, NHLBI TOPMed Program, Carole Sztalryd-Woodle, Braxton D. Mitchell, Matthew Healy, Elizabeth Streeten, Norann A. Zaghloul, Simeon I. Taylor, Jeffrey R. O’Connell, Alan R. Shuldiner

## Abstract

Increased LDL-cholesterol (LDL-C) and fibrinogen are independent risk factors for cardiovascular disease (CVD). We identified novel associations between an Amish-enriched missense variant (p.Asn352Ser) in a functional domain of beta-1,4-galactosyltransferase 1 (*B4GALT1*) and 13.5 mg/dl lower LDL-C (p=1.6E-15), and 26 mg/dl lower plasma fibrinogen (p= 9.8E-05). N-linked glycan profiling found p.Asn352Ser to be associated (p-values from 1.4E-06 to 1.0E-17) with decreased glycosylation of glycoproteins including: fibrinogen, ApoB100, immunoglobulin G (IgG), and transferrin. *In vitro* assays found that the mutant (352Ser) protein had 50% lower galactosyltransferase activity compared to wild type (352Asn) protein. Knockdown of *b4galt1* in zebrafish embryos resulted in significantly lower LDL-C compared to control, which was fully rescued by co-expression of 352Asn human *B4GALT1* mRNA but only partially rescued by co-expression of 352Ser human *B4GALT1* mRNA. Our findings establish *B4GALT1* as a novel gene associated with lower LDL-C and fibrinogen and suggest that targeted modulation of protein glycosylation may represent a therapeutic approach to decrease CVD risk.

---

Cardiovascular disease (CVD) accounts for 1 of every 3 deaths in the USA and is the leading cause of morbidity and mortality worldwide^1^. Elevated low-density lipoprotein cholesterol (LDL-C) increases arterial plaque formation and atherosclerosis, and fibrinogen increases risk for blood clotting and thrombosis; both are independent risk factors for coronary artery disease (CAD)^2,3^. LDL-C and fibrinogen are governed by both genetic and environmental factors, as well as the interplay between them^4,5^. Rare variants in *LDLR, PCSK9, APOB* cause severe forms of familial hypercholesterolemia, and many common genetic variants, each with small effect size, have been identified in large genome-wide association studies of both LDL-C and fibrinogen ^6-8^. However, few variants have been found with pleiotropic effects on more than one CAD risk factor. Similarly therapeutic approaches to mitigate CAD risk have focused on treating individual risk factors. Deeper understanding of the genetic determinants of LDL-C and fibrinogen may unveil novel targets for therapy that may be more efficacious and safe to treat or prevent CAD.

Founder populations can facilitate the identification of novel disease associations with variants that are enriched to a higher frequency through genetic drift. Multiple examples have been recently reported of highly enriched variants with large effect sizes associated with complex diseases and traits in homogenous populations in Iceland^9^, Sardinia^10^, Greenland^11^, Samoa^12^ and the Old Order Amish (OOA) ^13-19^. While such drifted variants are often rare or absent in the general population, their novel associations can inform biological mechanisms and therapeutic targets relevant to *all* humans.

The ability to perform whole-exome and whole-genome sequencing in founder populations provides the opportunity to identify drifted causative genetic variants hard to identify in general populations. In this study, we performed genetic association analyses for LDL-C in OOA participants with whole-genome and whole-exome sequencing data. We identified a novel genome-wide significant association between an Amish-enriched missense variant in the beta-1,4-galactosyltransferase 1 gene (*B4GALT1* p.Asn352Ser) and decreased LDL-C, which was also associated with decreased fibrinogen. Homozygotes for 352Ser have increased levels of incompletely synthesized glycans on glycoproteins as a result of decreased B4GALT1 enzymatic activity suggesting a potential mechanism to modulate lipid metabolism and coagulation.

## Association analyses identify *B4GALT1* p.Asn352Ser as a novel LDL-C variant

To identify genetic variants associated with LDL-C, we performed an exome-wide association scan using 5,890 OOA subjects with whole-exome sequencing (WES) performed at the Regeneron Genetics Center. Demographic and clinical characteristics of the WES subjects are shown in Supplementary Table 1. Linear mixed model association analysis identified several previously known loci for LDL-C as well as a novel locus on the short arm of chromosome 9 (Supplementary Fig. 1 and Supplementary Table 2). As shown in Figure 1a, a missense variant (rs551564683, p.Asn352Ser) in *B4GALT1* was strongly associated with 13.5 mg/dl lower LDL-C in an additive genetic model (p =1.6E-15). This variant has a minor allele frequency (MAF) of 6% in the OOA population while extremely rare in the general population (only 8 copies were identified in 140,000 WGS of non-Amish participants in the NHLBI Trans-Omics for Precision Medicine (TOPMed) program (www.nhlbiwgs.org).

**Fig. 1:**
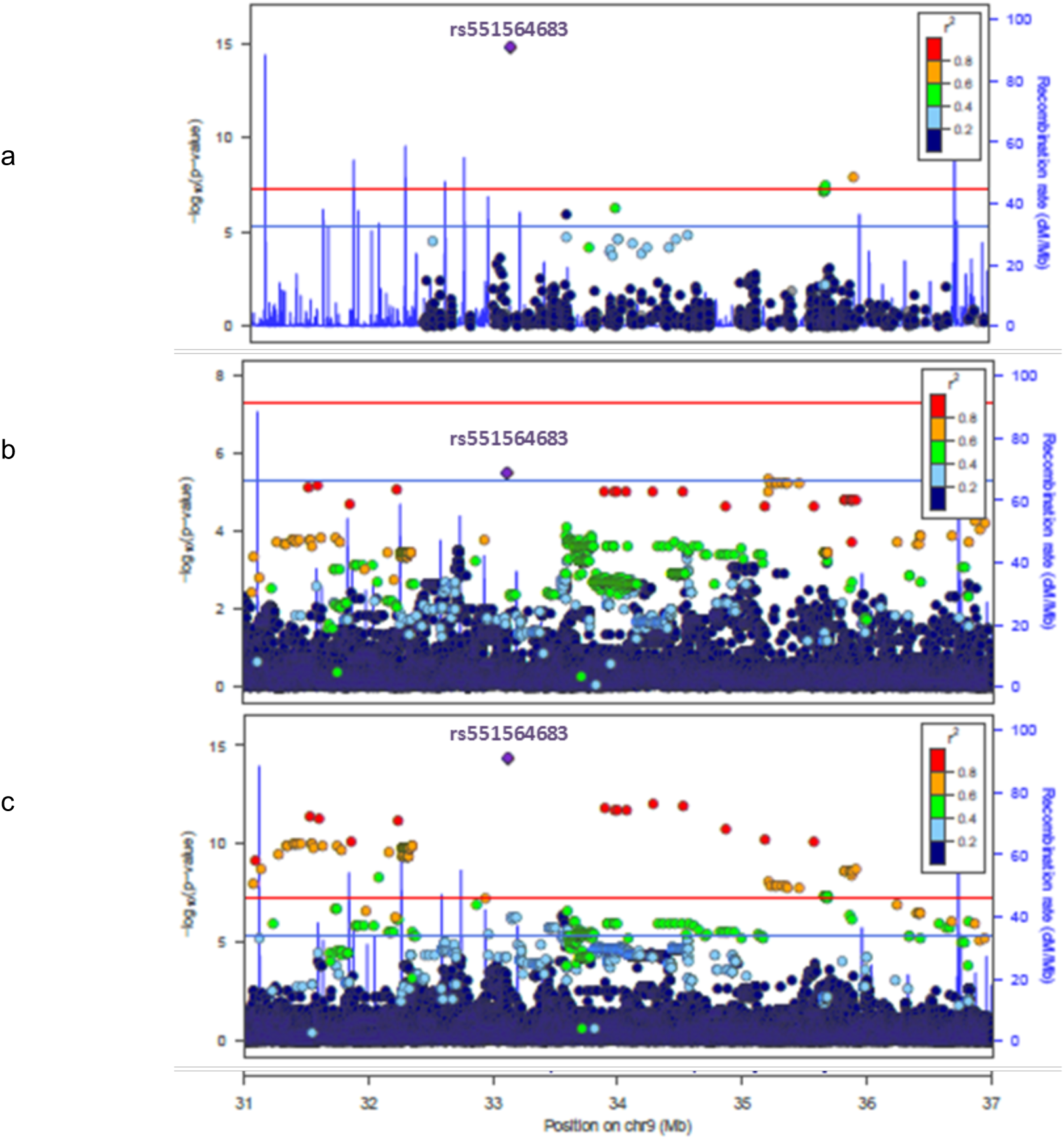
Evidence for genetic association between rs551564683, encoding *B4GALT1* p.Asn352Ser and LDL-C. **a**, Whole-exome sequencing results for 5,890 OOA subjects. **b**, Whole-genome sequencing results for 1,083 OOA subjects. **c**, Imputed data from genome-wide genotyping chip results for 5,890 OOA subjects. Blue line marks a genome-wide suggestive threshold (5.0E-06) and red line a genome-wide significant threshold (5.0E-08).

Since WES captures only the coding variants, we performed association analysis using 1,083 OOA subjects with whole-genome sequence (WGS) as part of TOPMed (dbGaP accession number: phs000956) to exhaustively interrogate all the coding and noncoding genetic variants in this region. Despite the much smaller sample size, the WGS analysis identified the same *B4GALT1* variant rs551564683 as the top signal in this region with p-value of 3.1E-06 and effect size of 16.9 mg/dl lower LDL-C (Figure 1b). In addition, WGS analysis revealed 20 variants in the region in high linkage disequilibrium (LD) with rs551564683 (r^2^ >0.8, range from 0.84 to 0.95), with LDL-C association p-values ranging from 6.3E-06 to 2.3E-05 (Supplementary Table 3). Of the 21 variants, rs551564683 is the only protein coding variant (*B4GALT1* p.Asn352Ser), and is classified as damaging or deleterious by 5 *in silico* protein function prediction algorithms (SIFT = deleterious, Polyphen2 = possibly damaging, LRT = deleterious, Mutation Taster = disease-causing, PROVEAN = deleterious). These 20 variants and rs551564683 comprise a 4 Mb Amish-specific haplotype extending from approximately 31.5Mb to 35.5Mb on chromosome 9 (Supplementary Fig. 2). These variants have a MAF of approximately 6% in the OOA while extremely rare or nonexistent in the general population (Supplementary Table 3)

**Fig. 2:**
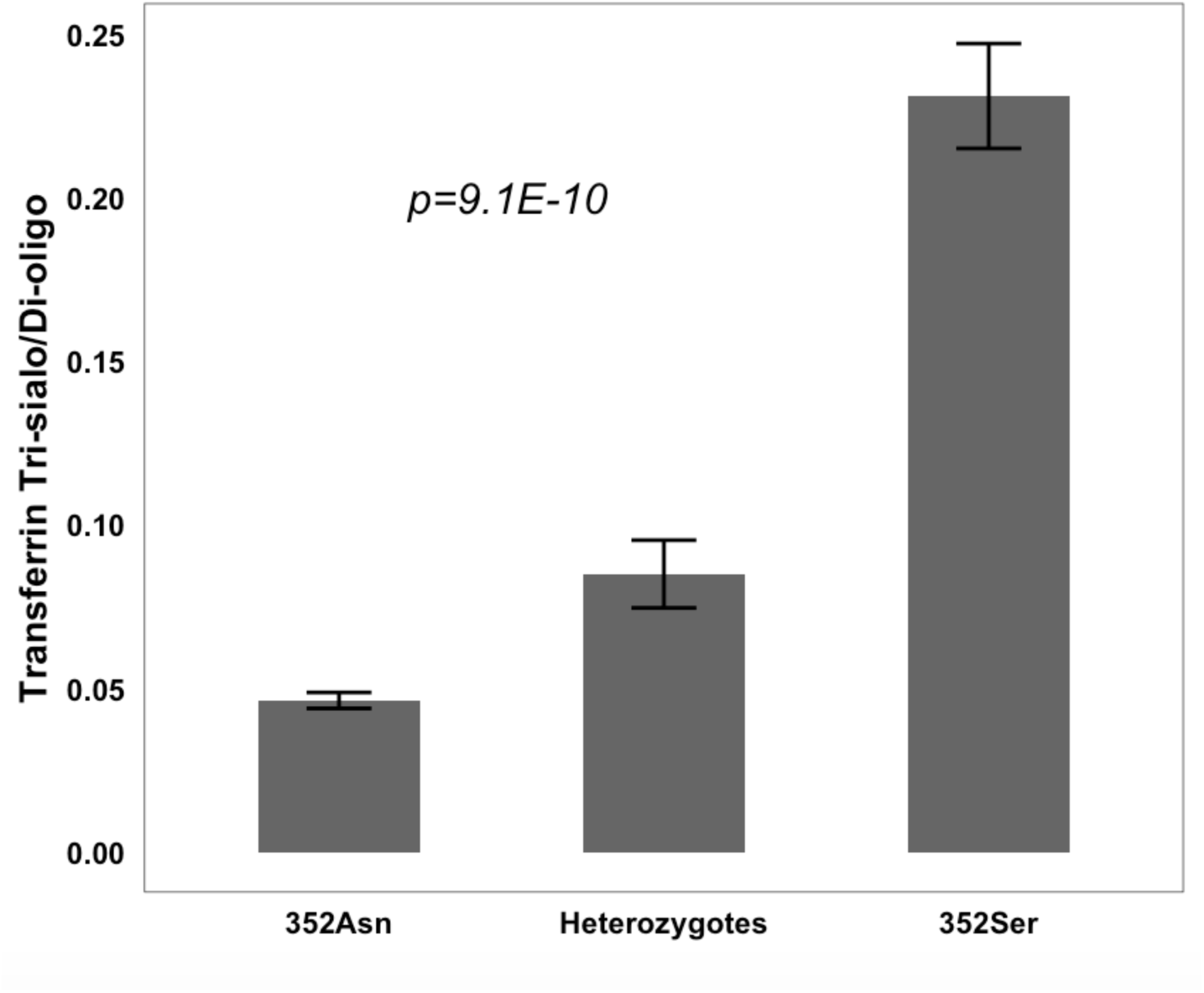
*B4GALT1* p.Asn352Ser is associated with increased carbohydrate-deficient transferrin. The transferrin Tri-sialo/Di-oligo ratio for the 3 genotype groups (11 352Asn homozygotes, 8 Asn352Ser heterozygotes and 9 352Ser homozygotes) as measured by the Carbohydrate Deficient Transferrin test.

The limited sample size of the WGS (n=1,083) was not able to differentiate statistically the top missense variant (p =3.1E-06) from the 20 other highly correlated variants (p from 6.3E-06 to 2.3E-05). To further differentiate between these 21 highly linked variants, we imputed genotypes in the full OOA dataset of 5,890 subjects with genotype chip data to the TOPMed WGS reference panel. As shown in Figure 1c and Supplementary Table 2, the missense variant was the top associated variant with a p value of 3.6E-15, two or more orders of magnitude smaller than any of the other variants (p from 9.4E-13 to 1.6E-09). Independent direct genotyping for 7 of these variants gave similar results (data not shown).

To determine if rs551564683 is the sole signal in the region, conditional analysis was performed. Conditional analysis adjusting for rs551564683 completely abolished the association of the other 20 variants while conditional analyses adjusting for any of the other 20 variants reduced the association of rs551564683 due to the strong correlation (r^2^ =0.84-0.95), but did not abolish it (p=1.0E-03 - 3.0E-07). Our analysis supports rs551564683 as the most likely causal variant in this region.

## Association with other traits and *B4GALT1* human knockout support a functional role of *B4GALT1* p.Asn352Ser

*B4GALT1* is a member of the beta-1,4-galactosyltransferase gene family which encode type II membrane-bound glycoprotein that plays a critical role in the processing of N-linked oligosaccharide moieties in glycoproteins. Impairment of B4GALT1 activity has the potential to alter the structure of N-linked oligosaccharides and introduce aberrations in glycan structure that have the potential to alter glycoprotein function.

A homozygous frame shift insertion in *B4GALT1* that results in a truncated dysfunctional protein was previously reported to cause Congenital Disorder of Glycosylation type 2 (CDGII) in 2 patients^20,21^. These 2 patients exhibited, among other traits, abnormal coagulation and very high levels of aspartate transaminase (AST). Interestingly we found *B4GALT1* p.Asn352Ser to be strongly associated with lower levels of fibrinogen (N= 805, beta = −26 mg/dl, p= 9.8E-05). We also found that the mean level of AST in 352Ser homozygotes was two-fold higher than that of 352Asn homozygotes (35.8 vs 18.5 U/L, respectively) (N= 5,595, additive model p= 2.3E-11, recessive model p= 2.4E-25), further supporting the functional role of this variant. While the high level of AST might suggest liver injury, we found no association with alanine aminotransferase (ALT), alkaline phosphatase (ALP), liver to spleen density ratio, or inflammatory markers (Supplementary Table 4). Moreover, all 352Ser homozygotes (N=12) had normal levels of gamma-glutamyl transpeptidase (GGT), activated partial thromboplastin time (aPTT), prothrombin time (PT), and internationalized normalized ratio (INR) (Supplementary Table 5).

## *B4GALT1* p.Asn352Ser is associated with increased levels of incompletely synthesized glycans on glycoproteins

To assess the impact of *B4GALT1* p.Asn352Ser on glycosylation, the carbohydrate deficient transferrin (CDT) test, which assesses glycosylation levels of transferrin and apolipoprotein CIII (ApoCIII), was performed using serum samples from 28 participants from the 3 genotype groups (9 352Ser homozygotes, 8 Asn352Ser heterozygotes and 11 352Asn homozygotes). As expected, based on the fact that *B4GALT1* plays a role in processing of N-linked carbohydrate, all samples had normal profiles for ApoCIII O-linked carbohydrate (Supplementary Table 6). All samples also had normal levels of mono-oligosaccharide/di-oligosaccharide transferrin ratio, and a-oligosaccharide/di-oligosaccharide transferrin ratio. However, while all eight 352Asn homozygotes had normal levels of the tri-sialo/di-oligosaccharide transferrin ratio, the ratio in all 352Ser homozygotes was abnormal; heterozygotes were intermediate between the two homozygote groups (p= 9.12 E-10) (Figure 2 and Supplementary Table 6). Since transferrin is mostly tetrasialylated^22,23^, this increase in trisialylated transferrin reflects a paucity of tetrasialylated molecules and indicates that the 352Ser allele is associated with increased levels of carbohydrate deficient transferrin, likely due to decreased enzymatic activity of B4GALT1.

To determine if the lower levels of sialylation are affecting only transferrin or extending to other glycoproteins, N-linked glycan profiling was performed using plasma samples from 24 participants (12 352Ser homozygotes and 12 352Asn homozygotes) for global plasma glycoproteins, as well as selected specific plasma glycoproteins - apolipoprotein B-100 (ApoB100), fibrinogen and immunoglobulin G (IgG) - by Hydrophilic Interaction Liquid Chromatography with Fluorescent Detection and Mass Spectroscopy (HILIC-FLR-MS).

Glycans from 352Ser homozygotes had increased levels of incompletely synthesized oligosaccharides as evidenced by increased percentages of truncated biantennary glycans devoid of galactoses and sialic acids (G0F, p=5.4E-11; bG0, p=2.1E-6), and biantennary glycans with only one galactose and one sialic acid (G1S1, p=5.1E-16). Reciprocally, 352Ser homozygotes had significantly lower levels of biantennary glycans with two galactose and two sialic acid residues (G2S2, p=2.6E-8) (Supplementary Table 7). The results for plasma ApoB100, fibrinogen and IgG were similar, where serum from 352Ser homozygotes had significantly increased level of glycans lacking galactose and sialic acid moieties and decreased levels of more mature glycans (Supplementary Table 8, 9, 10).

Overall, there was significantly lower galactosylation (p from 1.8E-08 to 1.0E-17) and sialylation (p from 1.4E-06 to 5.6E-15) for global glycoproteins, and enriched ApoB100, fibrinogen and IgG among 352Ser homozygotes compared to 352Asn homozygotes, however there was no difference in fucosylation (Figure 3), which is consistent with the role of B4GALT1 in adding galactose moieties that then are capped by sialic acid while having no role in the addition of fucose^24^.

**Fig. 3:**
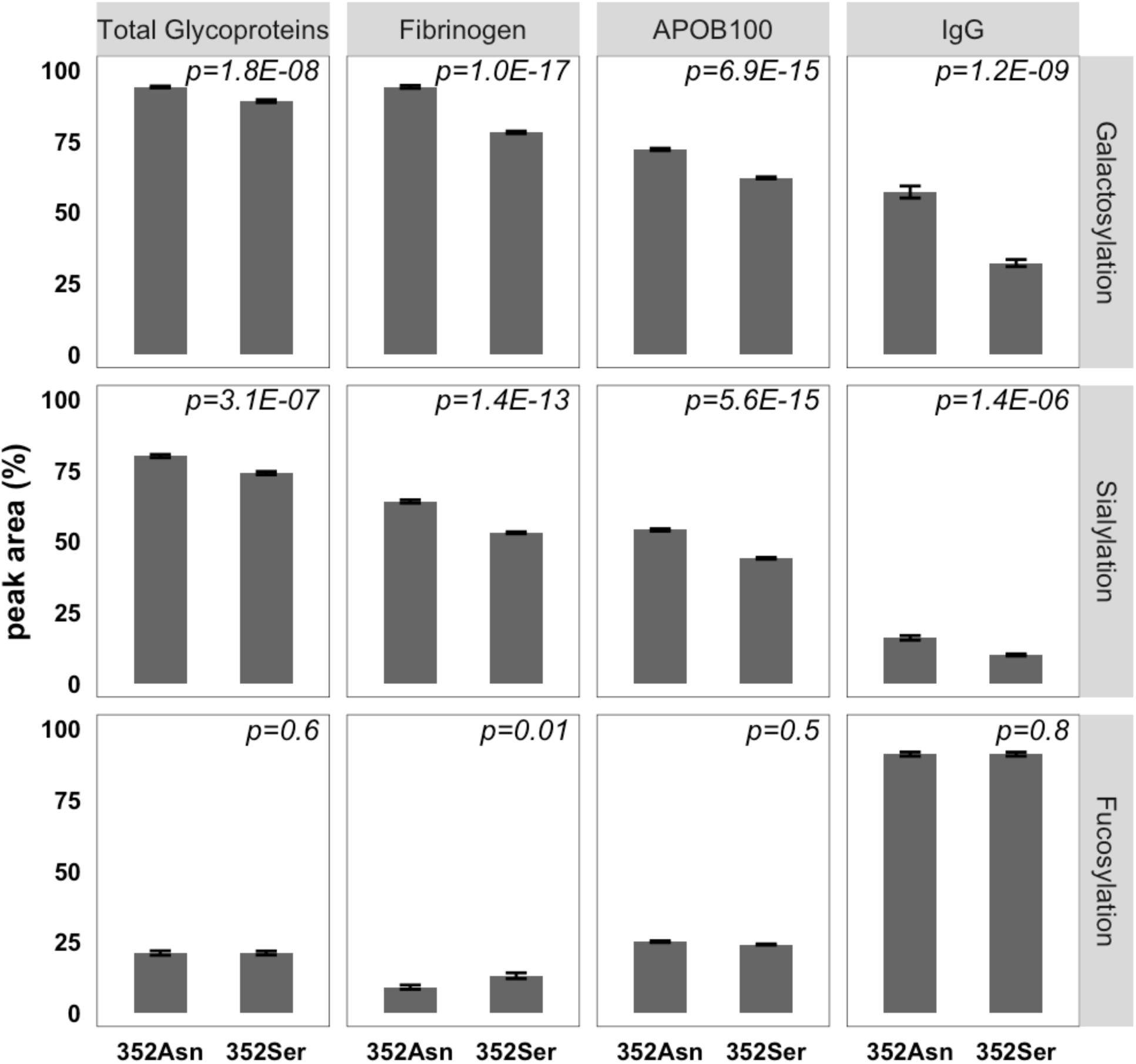
*B4GALT1* p.Asn352Ser is associated with decreased glycosylation. The levels of total galactosylation, sialylation and fucosylation in global plasma glycoproteins, fibrinogen, ApoB100, and IgG for 12 352Asn homozygotes and 12 352Ser homozygotes. Data are represented as mean and standard error.

In summary, the 352Ser allele is associated with significantly increased levels of incompletely synthesized glycans on glycoproteins indicating defective protein glycosylation.

## *B4GALT1* p.Asn352Ser causes reduced enzymatic activity

To compare the *in vitro* enzymatic activities of wild type (352Asn) and mutant (352Ser) B4GALT1 protein, we transiently transfected COS-7 cells, which express very low endogenous levels of B4GALT1, with cDNAs encoding myc-FLAG epitope-tagged versions of the proteins. We immunoprecipitated with anti-FLAG antibody, and measured B4GALT1 activities in the immune complexes (Supplementary Fig. 3). Compared to 352Asn B4GALT1, 352Ser B4GALT1 showed on average 50% decrease in galactosyltransferase specific activity (Figure 4a).

**Fig. 4:**
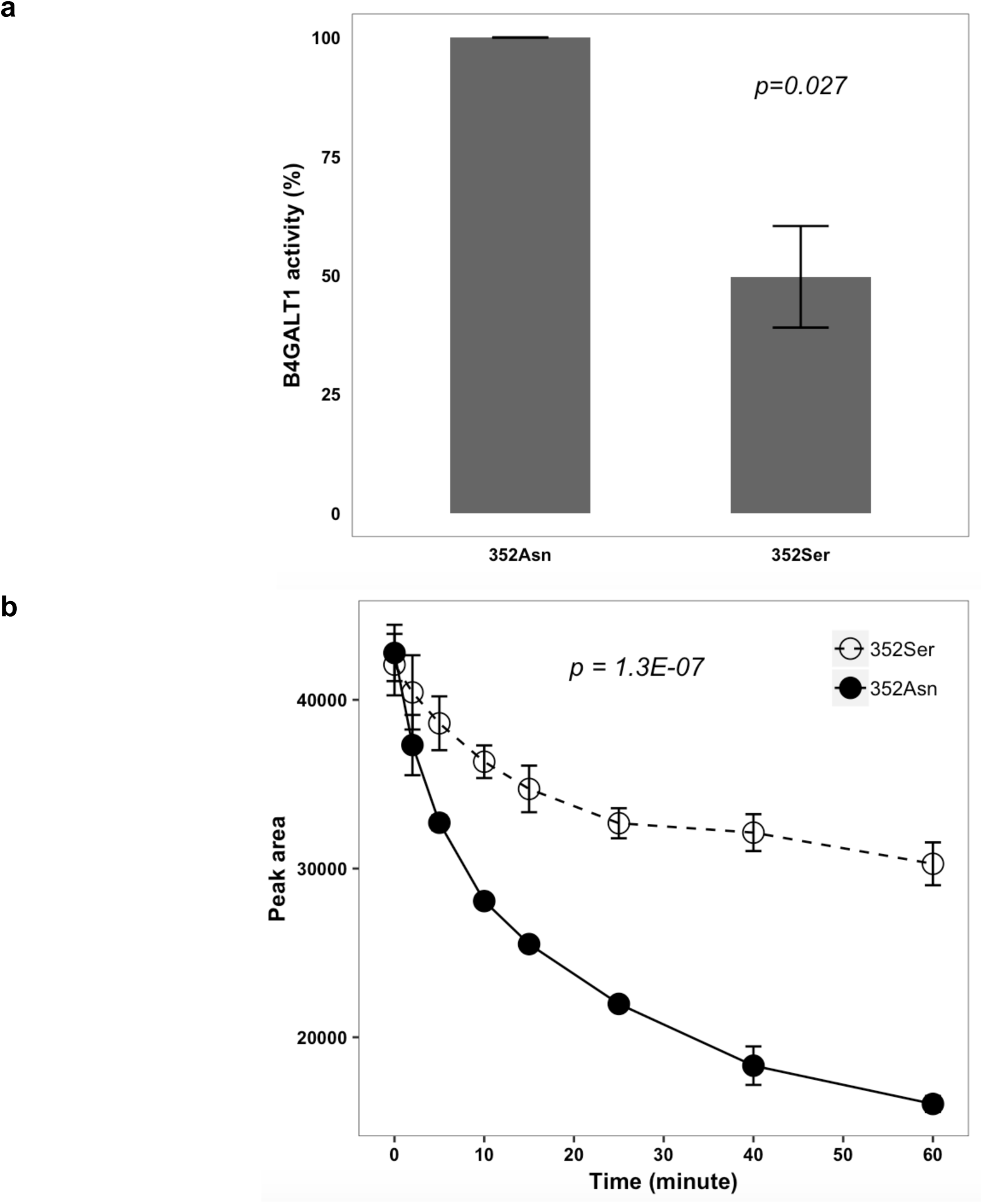
*B4GALT1* p.Asn352Ser decreases galactosyltransferase activity. **a**, The activity level of 352Asn B4GALT1 and 352Ser B4GALT1 immunoprecipitated proteins from transfected COS-7 cells. The data are expressed as the percent of 352Asn B4GALT1 activity and represent the mean of four experiments ± the standard error. **b**, The decrease in UDP-Gal incubated with recombinant B4GALT1 352Ser (open circles) and 352Asn proteins (solid circles).

As a complementary approach, we used synthesized recombinant human 352Asn and 352Ser B4GALT1 protein to test the reduction rate of the substrate uridine diphosphate galactose (UDP-Gal) incubated with each protein and found that the UDP-Gal level decreased faster with 352Asn than 352Ser B4GALT1, indicating decreased enzymatic activity of 352Ser B4GALT1 compared to 352Asn B4GALT1 (Figure 4b).

## Functional validation of the effect of *B4GALT1* p.Asn352Ser on LDL-C in zebrafish

We used a zebrafish model to investigate the effect of *B4GALT1* p.Asn352Ser on LDL-C using our established assays^19,25^. We first generated a genomic knockout of the zebrafish ortholog (*b4galt1*) using CRISPR/Cas9-mediated targeting of exon 2. Consistent with mouse reports of embryonic lethality in knockout animals^26^ injected F0 animals were not viable to adulthood and consistently died at juvenile stages. To circumvent the lack of viability, we employed a knockdown approach using a previously reported splice-blocking antisense morpholino oligonucleotide (MO) injected into embryos^27^. After validation of MO efficacy (Supplementary Fig. 4) and ruling out the possibility of off-target toxicity by demonstrating no increases in *d113 p53* expression and no overt morphological defects (Supplementary Fig. 5, 6a), we quantified changes in LDL-C levels in unfed larvae at 5 days post fertilization (dpf) as per previously published protocols^25^. We observed a significant decrease in LDL-C in MO-injected larvae compared to control larvae, consistent with a role for *b4galt1* in LDL-C homeostasis (Figure 5). We further confirmed this result using a second splice-blocking MO targeting exon 2 of *b4galt1* which similarly produced a reduction in LDL-C concentration (Supplementary Fig. 6b-d).

**Fig. 5:**
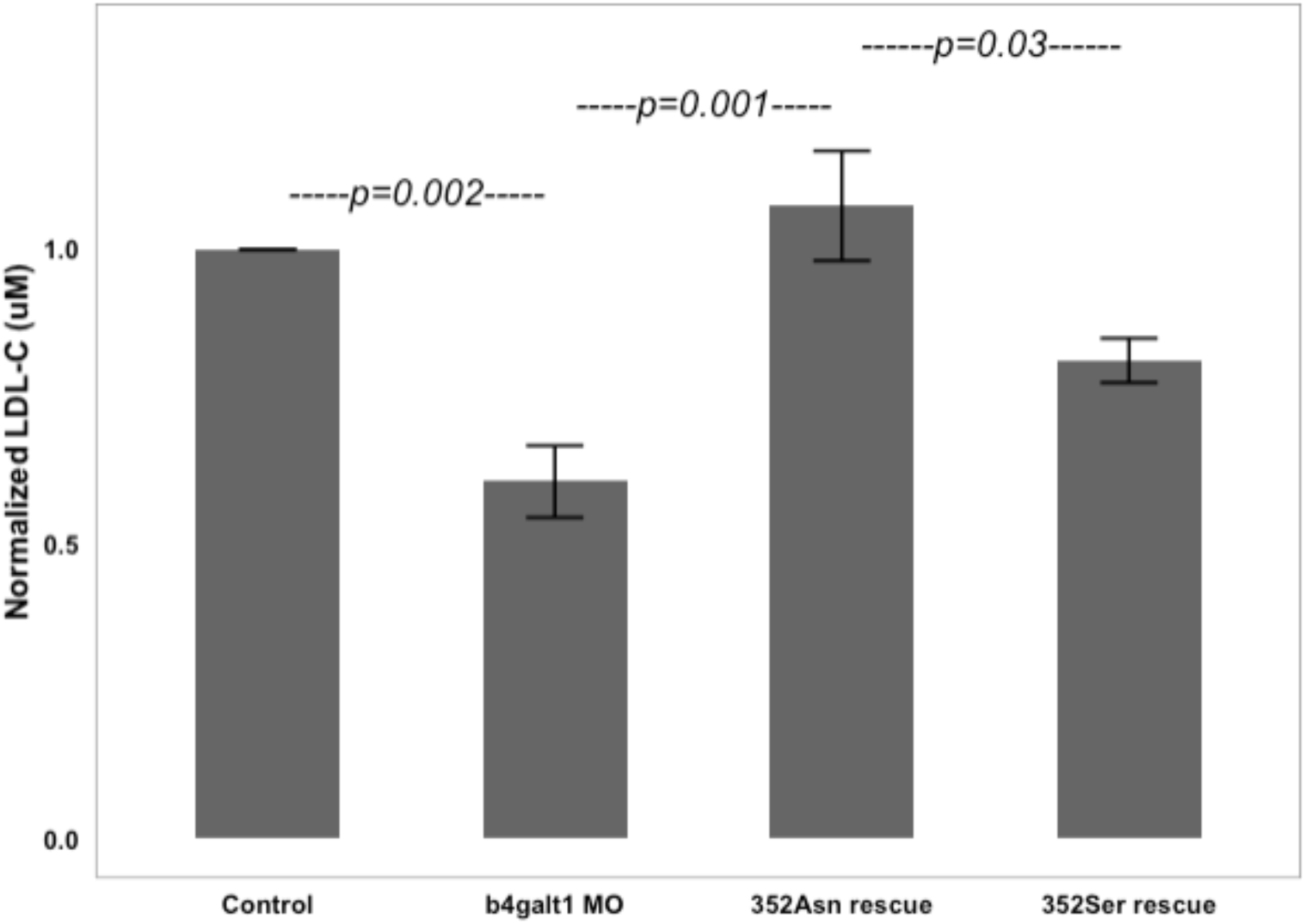
Quantification of LDL-C in zebrafish. Average LDL-C in homogenates of unfed wild type 5 dpf zebrafish larvae (n=50 per well, duplicated within each experiment, n=7 for Control, and 352Asn rescue, n=5 for b4galt1 MO, n=4 for 352Ser rescue). Larvae were first injected with either a non-targeting control MO (Control) or 8 ng of morpholino (b4galt1 MO) against b4galt1 at 1-2 cell stage. Rescue conditions were 8 ng MO co-injected with 100pg 352Asn human B4GALT1 mRNA (352Asn rescue), or 8 ng MO co-injected with 100 pg B4GALT1 mRNA encoding the 352Ser mutation (352Ser rescue). Data are represented as mean and standard error, normalized to control MO within each experiment and averaged across experiments.

To validate the specificity of these observations and to test the functionality of human *B4GALT1* in zebrafish, we co-injected mRNA encoding human 352Asn *B4GALT1* along with *b4galt1* MO into embryos and assessed LDL-C in unfed larvae. This resulted in LDL-C levels that were statistically indistinguishable from those in larvae injected only with a control MO (p=0.45), suggesting that the human mRNA could rescue the effects of knockdown of the zebrafish gene (Figure 5).

These data suggest that human wild type *B4GALT1* mRNA is functional in zebrafish and support the use of this model system for functional interpretation of p.Asn352Ser. Using site-directed mutagenesis^28^, we introduced a T to C change in the coding sequence of human *B4GALT1* and generated full length mRNA. Co-injection of the 352Ser *B4GALT1* mRNA with *b4galt1* MO resulted in a reduced capacity for rescue of the LDL-C phenotype. The resulting LDL-C concentration was 25% lower than that resulting from co-injection of 352Asn *B4GALT1* mRNA with *b4galt1* MO, a statistically significant effect (p=0.03). This level of LDL-C was also statistically greater, however, than *b4galt1* MO alone (p=0.02) (Figure 5), suggesting only a partial defect in function introduced by the missense variant.

To examine the relevance of *b4galt1* to other hypercholesterolemic models, we examined the impact of *b4galt1* knock down on two previously reported zebrafish models of elevated LDL-C: high cholesterol diet fed larvae and *ldlr* knockdown animals^25^. For the former, we treated 5 dpf larvae with either a control diet or the same diet supplemented with 4% w/w cholesterol as previously reported^25^. While *b4galt1* knockdown reduced LDL-C in animals fed a control diet, this effect was abolished with introduction of a high cholesterol diet (Supplementary Fig. 7). In contrast, when we introduced *b4galt1* suppression on a background of genetically-induced hypercholesterolemia via *ldlr* MO injection, we observed amelioration of the LDL-C phenotype in those animals (Supplementary Fig. 7). These data suggest that suppression of *b4galt1* can abrogate elevated LDL-C resulting from genetic defects in LDL-C metabolism but not that introduced by diet.

## Discussion

Large genome-wide analyses of approximately 600,000 individuals have identified 386 loci associated with lipid traits, none of which identified the *B4GALT1* gene^29^. Using next generation sequencing in 5,890 OOA, we found a strong novel association of *B4GALT1* p.Asn352Ser with lower plasma LDL-C and fibrinogen levels. Glycosylation profiling and experimental assays confirmed the functional role of this variant. This variant is very rare in the general population, but has 6% frequency in the Amish, likely due to genetic drift over approximately 14 generations after founding. This report highlights the value of founder populations in identifying novel gene variants that can provide new insights into human biology of common traits.

Homozygosity for a protein truncating mutation in *B4GALT1* (1032insC) reported in two related patients is known to cause CDG type 2 (CDGII)^20,21^. Both patients have marked abnormal carbohydrate structures on glycosylated proteins and severe clinical phenotypes manifesting in early childhood with developmental delay, hypotonia, coagulopathy, and elevated transaminases^20,21,30^. Knock-out of *b4galt1* in mice results in semi-lethality after birth and several other sever developmental abnormalities ^31,32^. Interestingly, mass spectrometry studies of N-glycans from plasma and hepatic membrane glycoproteins, revealed an unexpected high level of sialylation and galactosylation in b4galt1 -/-(∼60% compared to that of b4galt1+/+ mice). Kotani *et. al*., explained the altered pattern of sialylation and galactosylation with a shift in synthesis from type 2 to type 1 glycan chains in *b4galt1* ^*-/-*^ versus *b4galt1*^*+/+*^ mice, most probably caused by b3galt proteins in response to loss of *b4galt1*^33^. Conversely, the overall decrease in galactosylation and sialylation that we observed in the carriers of 352Ser *B4GALT1* missense variant, suggests that this missense variant could act as hypomorph. Indeed, our preliminary investigation showed that the level of type 1 chain (corresponding to the b 1,3 linkage) was very low and comparable between 352Asn and 352Ser homozygotes (data not shown), suggesting that our missense mutation did not cause any glycan chain type switch as shown in Kotani *et. al*., and that compensatory action of *B3GALT1* may differ between mouse and human.

The p.Asn352Ser *B4GALT1* missense variant that we identified in the Amish does not appear to be associated with any severe phenotype. In fact, the self-reported medical and family history of fifteen 352Ser homozygotes did not note previous heart attack, stroke, coronary angiogram, blocked arteries, or heart/carotid artery surgery. None had evidence of coronary ischemia or MI on EKG except for one 70 year old person who showed evidence of a potential previous MI. Abdominal ultrasound from five 352Ser homozygotes over 50 years old showed normal abdominal aorta with no atherosclerotic plaque, fatty liver or aneurysm. The variant showed no association with reduced coronary artery calcification (CAC) or aortic calcification (AC), however, the sample size was small with relatively young average age of 52, that included only one 352Ser homozygote (Supplementary Table 4).

We also found that 352Ser homozygotes had double the serum AST levels compared to 352Asn homozygotes, however, they all had normal levels of ALT, GGT and ALP, suggesting that the source of high AST is from tissue other than liver, *i*.*e*., muscle. Moreover, they also had normal INR, PTT and inflammation markers (Supplementary Tables 4 and 5). The absence of a severe phenotype in the minor homozygotes may be due to the subtle effect of the missense mutation compared to the protein truncating frameshift mutation identified in CDGII patients.

The missense variant p.Asn352Ser changes the asparagine to serine at position 352 in the 398 amino acid sequence of the long isoform of human *B4GALT1* (corresponding to position 311 of the short isoform). The asparagine residue is completely conserved across 100 vertebrate species (GERP score = 5.9). p.Asn352Ser is located in the highly flexible long C terminal region of the protein that undergoes conformational changes to allow for the exchange of the sugar molecule during glycosylation^34^. Hence, a mutation in this region may impede the necessary conformational change and impact glycosylation efficiency as shown by our glycosylation profiling and functional assays. However, it is important to note that the 352Ser mutation had less severe effects on glycosylation than that previously reported in CDGII patients carrying a *B4GALT1* truncating mutation^20^.

The mechanism by which this variant leads to lower LDL-C and fibrinogen remains to be elucidated. B4GALT1 is a ubiquitously expressed protein that transfers the galactose from uridine diphosphate galactose (UDP-Gal) to specific glycoprotein substrates^21^. A defect in *B4GALT1* that affects glycosylation and sialylation may affect the folding, secretion, stability, activity, and half-life of lipoprotein and coagulation related glycoproteins^35-37^ ultimately resulting in lower levels of circulating LDL-C and fibrinogen.

Genome-wide association studies (GWAS) have identified common non-coding variants at the *B4GALT1* locus that are significantly associated with IgG glycosylation and AST levels^24,38^, but not with any lipid traits. However, several lines of evidence point to protein glycosylation as a major player in lipid metabolism. Reduced APOB glycosylation leads to shorter LDL-C half-life and rapid clearance from the circulation, which in turn reduces its atherogenic effect^35^, while increased LDL-C glycosylation was associated with increased LDL-C oxidation leading to greater atherosclerosis ^39^. Glucose mediated N-glycosylation plays a major role in regulating sterol regulatory element-binding proteins (SREBPs) which are major players in cholesterol metabolism^40^. Also, reduced LDLR glycosylation was found to result in reduced lipid binding and endocytosis^41^. Recently, changes in IgG glycosylation have been associated with blood lipids and dyslipidemia in Han Chinese^42^. A similar role for glycosylation was also reported in relation to coagulation proteins, where mutations affecting glycosylation were associated with different enzymatic activity^37^. Finally, GWAS has identified loci containing other enzymes involved in glycoprotein biology associated with lipid traits including *GALNT2*^43^, *ST3GAL4*^6^, *and ASGR1*^44^.

Similarly, studies support a role of glycosylation in CVD. For example, a loss of function mutation in *ASGR1* was reported to be associated with lower non-HDL cholesterol and reduced coronary artery disease (CAD) risk^44^. *ASGR1* encodes a subunit of the asialoglycoprotein receptor (the “Ashwell receptor”) that mediates binding and endocytosis of asialylated N-glycoproteins. Recently Menni *et*.*al*.^45^ reported that the glycosylation profile of IgG was associated with CVD risk score and subclinical atherosclerosis in two independent cohorts. While these studies are not specific to *B4GALT1*, they support an important role of glycosylation in CVD with potential prognostic and/or therapeutic implications.

In summary, we discovered a novel missense variant in *B4GALT1* that is associated with lower LDL-C and fibrinogen, both cardioprotective phenotypes, through enrichment in a founder population. Evidence from human data as well as *in vitro* and animal-based experiments indicate that the variant leads to decreased protein glycosylation. Further understanding of the underlying biological mechanism of the variant may provide new insights into the role of glycosylation in CVD risk that may lead to novel therapeutic targets.

## Online content

Methods, additional references, Nature Research reporting summaries, statements of data availability and associated accession codes are available in the online paper

## Acknowledgements

This work was supported in part from NIH grants U01 HL137181, U01 HL072515, R01 AG18728, R01 HL121007, P30 DK072488, AHA 17GRNT33661168 and Regeneron Pharmaceuticals, Inc.

Whole-genome sequencing (WGS) for the Trans-Omics in Precision Medicine (TOPMed) program was supported by the National Heart, Lung and Blood Institute (NHLBI). WGS for “NHLBI TOPMed: Genetics of Cardiometabolic Health in the Amish” (phs000956) was performed at the Broad Institute of MIT and Harvard (HHSN268201500014C). Centralized read mapping and genotype calling, along with variant quality metrics and filtering were provided by the TOPMed Informatics Research Center (3R01HL-117626-02S1). Phenotype harmonization, data management, sample-identity QC, and general study coordination, were provided by the TOPMed Data Coordinating Center (3R01HL-120393-02S1).

We gratefully thank the Amish community and research volunteers for their long-standing partnership in research, and acknowledge the dedication of the Amish liaisons, field workers and the Amish Research Clinic staff, without which these studies would not have been possible.

## Author contributions

Conceived, designed and supervised the work: MEM, ARS, ANE

Performed data collection: MEM, ARS, JO, BS, NL, GDG, GT, LM, AR, RMc, NAZ, WL, YM

Contributed to data analysis MEM, CVH, BS, NL, BY, GDG, GT, LM, MH, JRO, RMc, NAZ, KAR, JP

Interpreted the results: MEM, ARS, CVH, BS, NL, TJD, GDG, ANE, MH, JRO, AH, CS, BDM, ES, NAZ, SIT

Preparation of the manuscript: MEM, ARS

