## Supplemental Figures and Tables for "Genetic and functional evidence relates a missense variant in *B4GALT1* to lower LDL-C and fibrinogen"

[4]Full list of names and affiliations in the online supplementary materials

[5]Geriatrics Research and Education Clinical Center, Baltimore VA Medical Center, Baltimore, MD 21201, USA

### TABLE OF CONTENTS

#### Supplementary Figures

**Supplementary Figure 1**: Exome-wide association results for LDL-C **3-4**

**Supplementary Figure 2:** The 4 Mb Amish-specific haplotype on chr9p21.1 **5**

**Supplementary Figure 3:** The effect of the p.Asn352Ser mutation on galactosyltransferase activity **6**

**Supplementary Figure 4:** Reduced *b4galt1* mRNA in 5 dpf zebrafish larvae detected by qRT-PCR **7**

**Supplementary Figure 5:** Diagnostic marker (delta113 isoform of *p53* gene) of MO-induced off-target in 5 dpf zebrafish larvae. **8**

**Supplementary Figure 6:** Validation of MO efficacy and specificity**.** **9**

**Supplementary Figure 7:** b4galt1 knockdown in hypercholesterolemic conditions **10**

#### Supplementary Tables

**Supplementary Table 1:** Demographic and clinical characteristics of the OOA study population. **11**

**Supplementary Table 2:** Top associated variants with LDL-C from whole exome sequencing **12**

**Supplementary Table 3:** Top associated variants with LDL-C at the *B4G[illegible]* locus from whole genome sequencing. **13**

**Supplementary Table 4:** The association between *B4GALT1* p.Asn352Ser and relevant traits in the OOA. **14**

**Supplementary Table 5:** The ranges of GGT and coagulation measures in 12 subjects with *B4GALT1* 352Ser homozygotes. **14**

**Supplementary Table 6:** Carbohydrate deficient transferrin test results **15**

**Supplementary Table 7:** peak area of significantly different glycans in plasma N-linked glycoproteins **15**

**Supplementary Table 8:** peak area of significantly different glycans in plasma N-linked APOB100 **15**

**Supplementary Table 9:** peak area of significantly different glycans in plasma N-linked Fibrinogen **16**

**Supplementary Table 10:** peak area of significantly different glycans in plasma N-linked IgG **16**

### Supplementary Figures

**a**

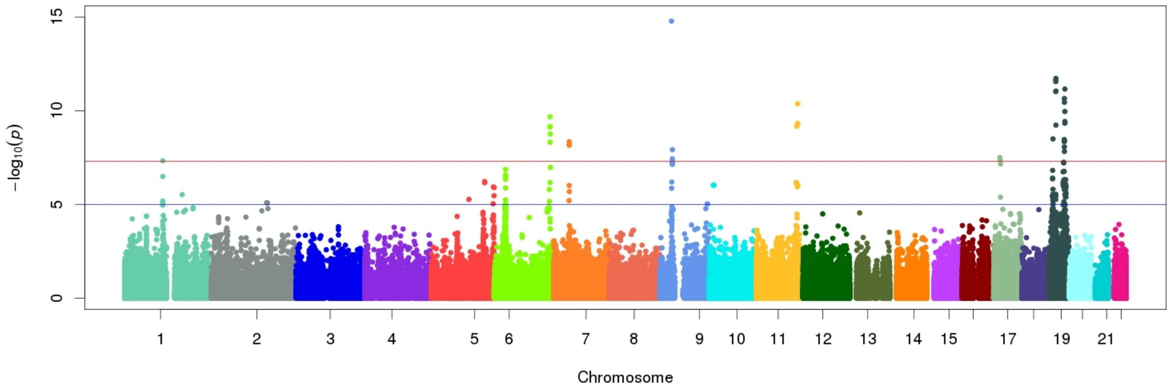

**b**

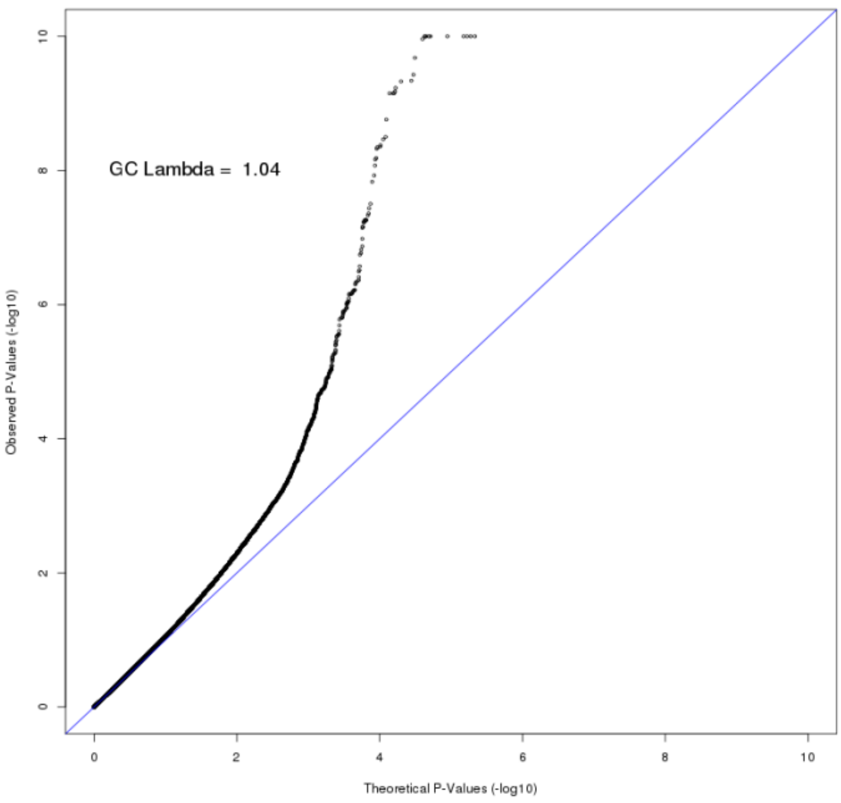

**c**

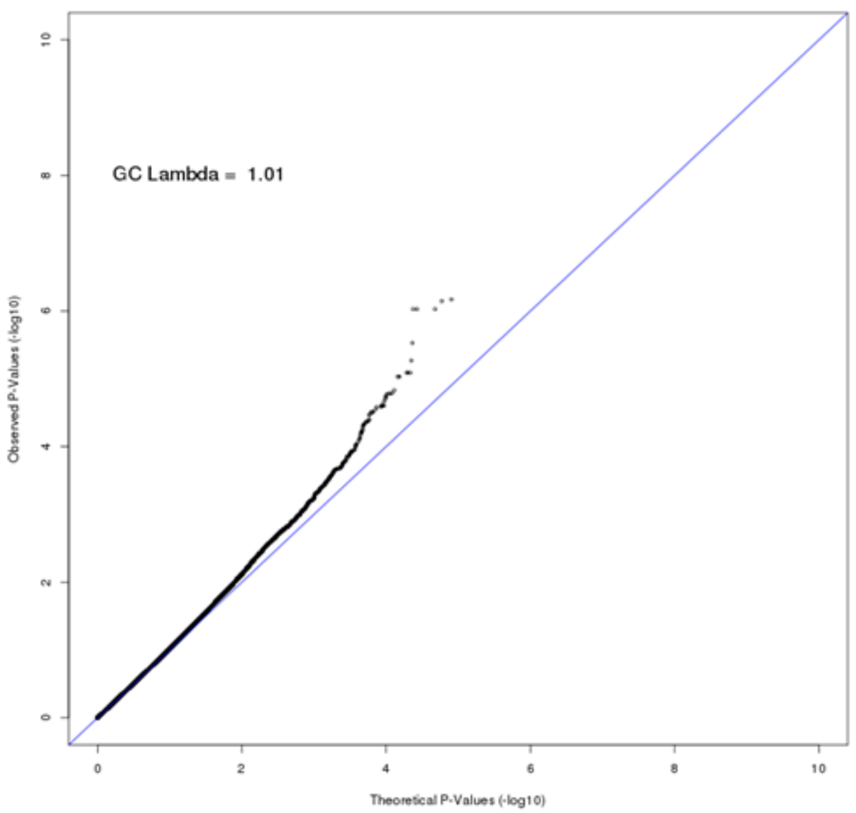

**Supplementary Fig. 1: Exome-wide association results for LDL-C. a,** Manhattan plot for the exome-wide association results of LDL-C in 5,890 OOA, adjusted for age, age squared, sex, Amish sub-study, familial correlation, and *APOB* p.Arg3527Gln.. **b,** QQ plot of association p-values of all results (GC Lambda =1.04). **c,** QQ plot of association p-values after removing SNPs within 10 Mb of each signal (GC Lambda =1.01). Blue line marks a genome-wide suggestive threshold (5.0E-06) and red

| Frequency | rs577483462 | rs539837022 | rs115335790 | rs544554676 | rs551564683 | 9:33892787 | rs1012682673 | rs998550643 | rs1023990285 | 9:34277757 | rs898800582 | rs1040328811 | rs536902364 | rs1054917306 | rs186441167 | rs370946679 | rs375489986 | rs118137738 | rs139585733 | rs569363002 | rs149557496 |
|---|---|---|---|---|---|---|---|---|---|---|---|---|---|---|---|---|---|---|---|---|---|
| **108** | 1 | 1 | 1 | 1 | 1 | 1 | 1 | 1 | 1 | 1 | 1 | 1 | 1 | 1 | 1 | 1 | 1 | 1 | 1 | 1 | 1 |
| **11** | 1 | 1 | 1 | 1 | 1 | 1 | 1 | 1 | 1 | 1 | 1 | 1 | 1 | 1 | 0 | 0 | 0 | 0 | 0 | 0 | 0 |
| **7** | 1 | 1 | 1 | 1 | 1 | 0 | 0 | 0 | 0 | 0 | 0 | 0 | 0 | 0 | 0 | 0 | 0 | 0 | 0 | 0 | 0 |
| **6** | 0 | 0 | 0 | 0 | 1 | 1 | 1 | 1 | 1 | 1 | 1 | 1 | 1 | 1 | 1 | 1 | 1 | 1 | 1 | 1 | 1 |

**Supplementary Fig. 2: The 4 Mb Amish-specific haplotype on chr9p21.1.** Frequency and haplotype structure of the top 21 LDL-C associated variants in 136 heterozygotes for p.Asn352Ser (rs55156483).

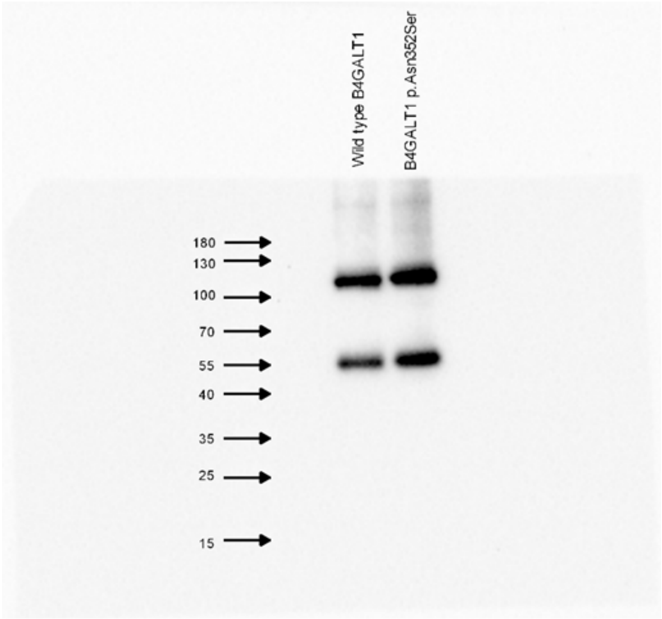

**Supplementary Fig. 3: The effect of the p.Asn352Ser mutation on galactosyltransferase activity.** Western blot of epitope-tagged wild type B4GALT1 and B4GALT1 352Ser proteins immunoprecipitated from COS-7 cells using Anti-FLAG M2 and detected using the anti-myc antibody. All of the immunoblot experiments demonstrated two bands: a 55 kDa band corresponding to the expected size of myc-Flag-B4GALT1 and a 110 kDa band. The B4GALT1 catalytic domain has previously been reported to form dimers[46], which raises the possibility that the 110 kDa band corresponds to B4GALT1 dimer. The previously reported B4GALT1 catalytic domain dimer was only detected in non-denaturing gels. In contrast, the 110 kDa band was demonstrated by SDS-polyacrylamide gel electrophoresis in the presence of beta-mercaptoethanol. Suggesting that the dimer is stabilized by a non-disulfide covalent bond. The abilities of the recombinant enzymes to transfer UDP from UDP-galactose to N-Acetyl-D-glucosamine were normalized to the densitometric sum of the 55 kDa and 110 kDa bands of eluted myc-B4GALT1 detected on western blots.

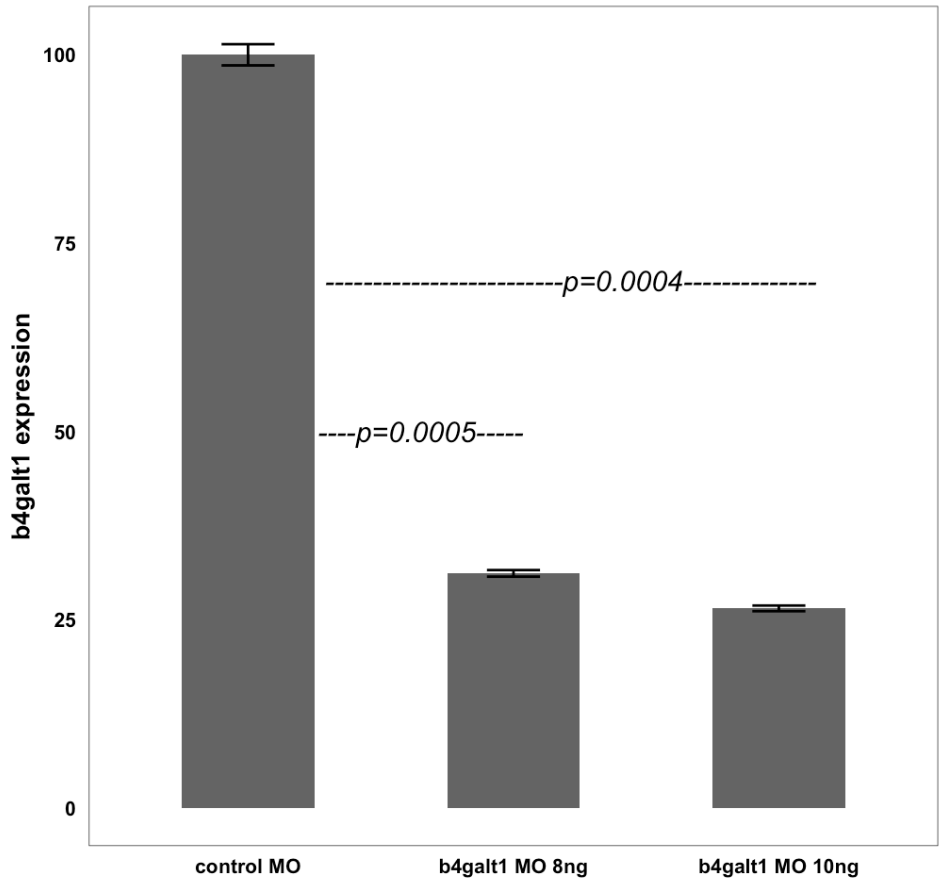

**Supplementary Fig. 4: Reduced *b4galt1* mRNA in 5 dpf zebrafish larvae detected by qRT-PCR.** Quantification used cDNA generated from RNA isolated from 5 dpf larvae injected with either control MO or indicated concentration of *b4galt1* MO (n=50 larvae per experiment, n=2 experiments, concentration shown relative to beta-actin and normalized to control MO). Data are represented as mean and standard error.

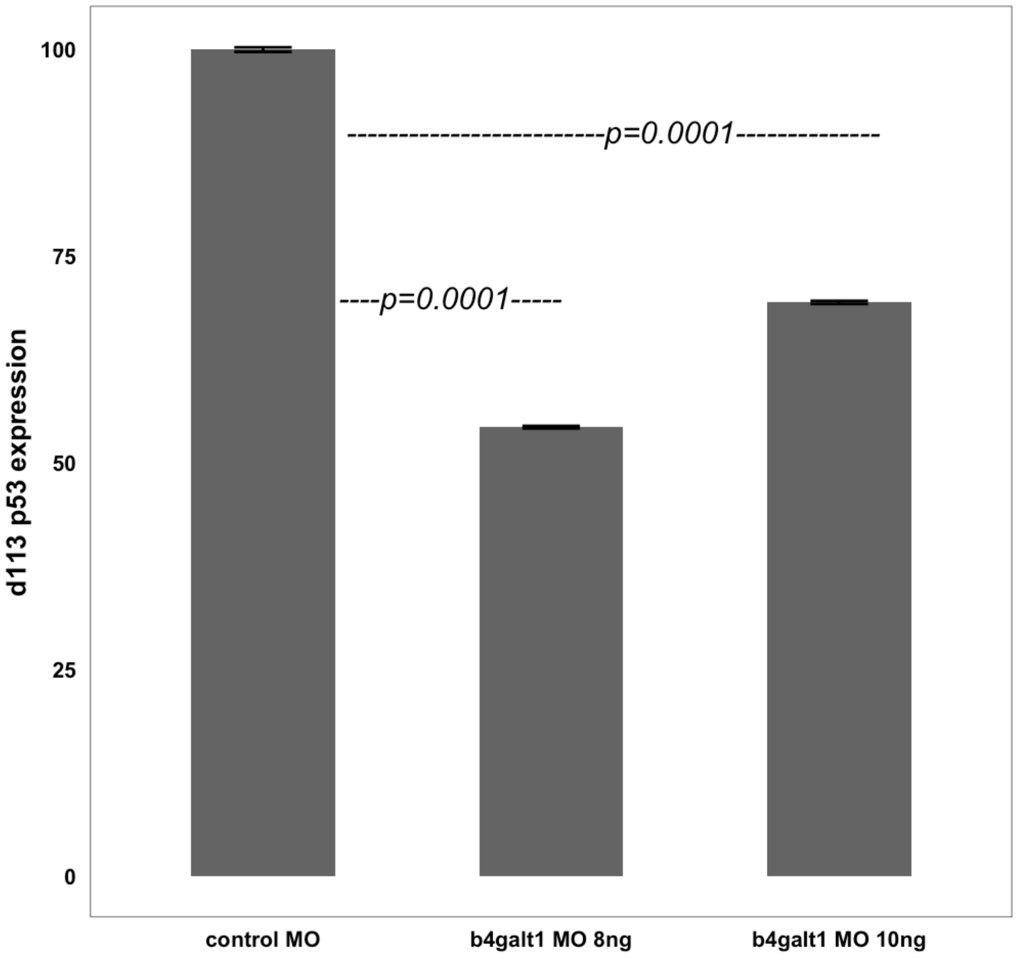

**Supplementary Fig. 5: Diagnostic marker (delta113 isoform of *p53* gene) of MO-induced off-target in 5 dpf zebrafish larvae.** qRT-PCR quantification using cDNA generated from RNA isolated from 5 dpf larvae injected with either control MO or *b4galt1* MO at the indicated concentration (n=50 larvae per experiment, n=2 experiments, concentration shown relative to beta-actin and normalized to control MO). Data are represented as mean and standard error.

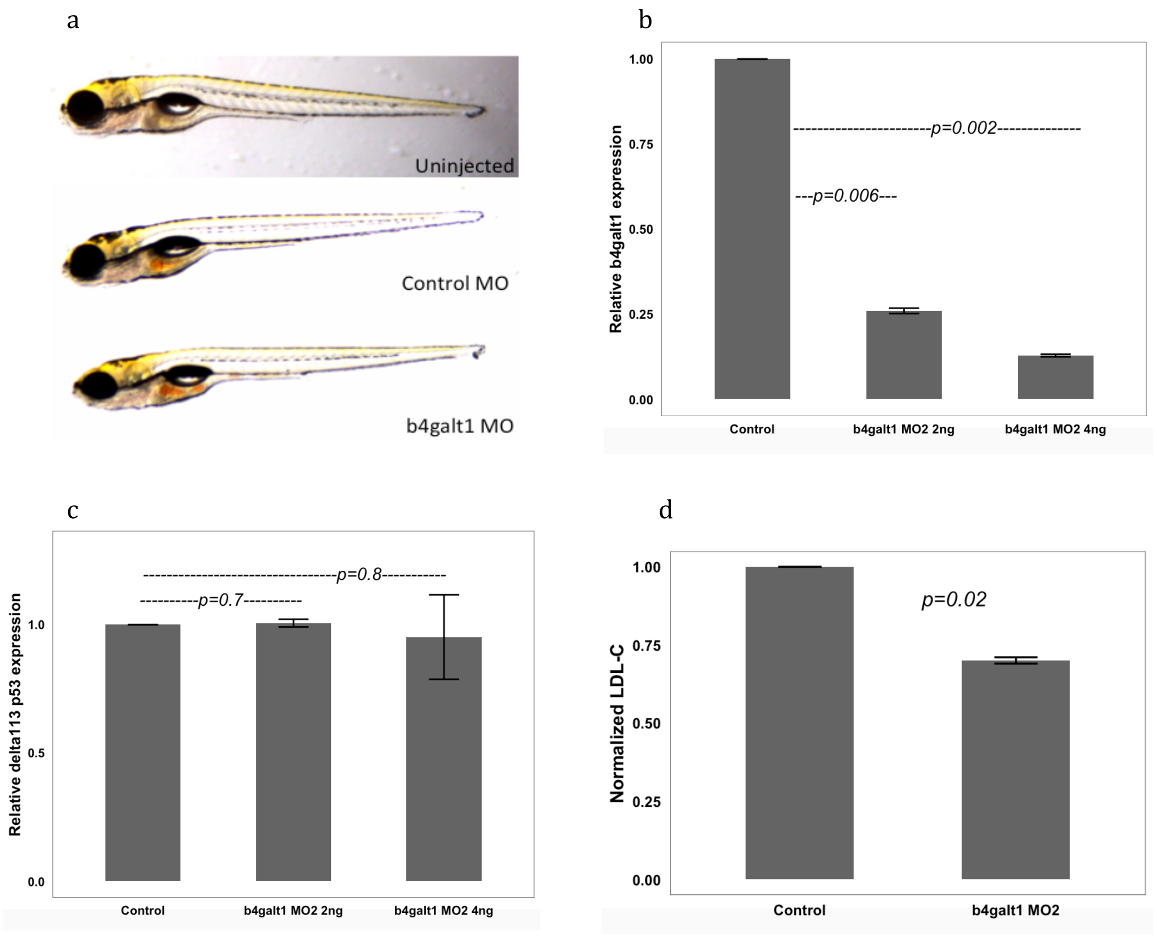

**Supplementary Fig. 6: Validation of MO efficacy and specificity. a**, Representative brightfield images of uninjected or indicated MO injected larvae at 5dpf. Scale bar=1mm. **b-c,** Quantification of relative b4galt1 RNA (**b**) or delta113 isoform of *p53* (**c**) expression in 5dpf larvae injected at the indicated concentrations with a second MO targeting *b4galt1* splicing (n=50 larvae per experiment, n=2 experiments, concentration shown relative to beta-actin and normalized to control MO). **d,** Average LDL-C, normalized to control MO, in 5 dpf larvae injected with the splice-blocking MO at 4ng concentration (n=50 larvae per experiment, n=3 experiments). All data are represented as mean and standard error.

**a**

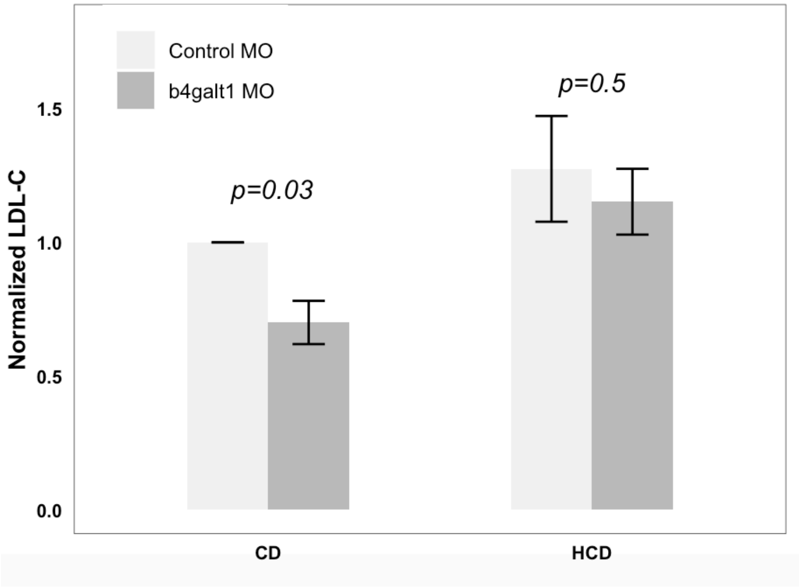

**b**

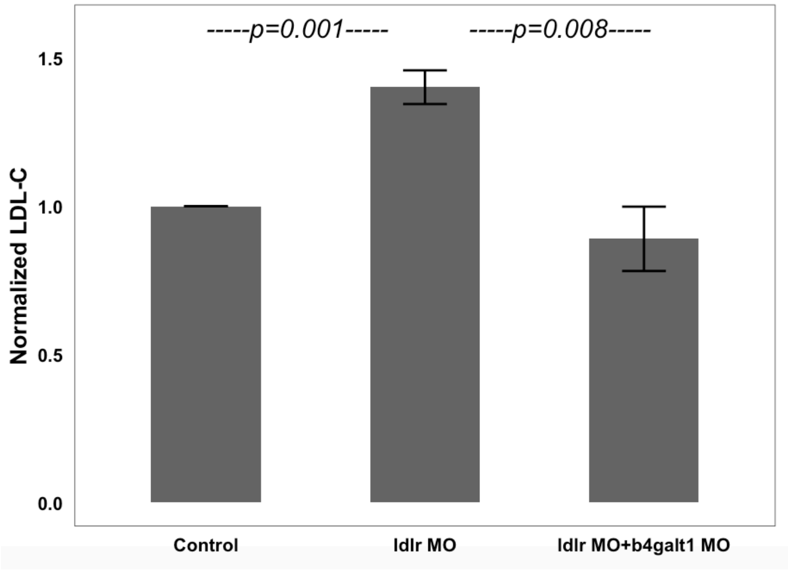

**Supplementary Fig. 7: b4galt1 knockdown in hypercholesterolemic conditions. a,** Quantification of average LDL-C in 6 dpf larvae injected with control MO or *b4galt1* MO and fed either a control diet (CD) or a high cholesterol diet (HCD), normalized to control MO on CD. (n=50 larvae per experiment, n=4 experiments). **b,** Average LDL-C in unfed 5 dpf larvae injected with indicated MOs, normalized to control MO (n=50 larvae per experiment, n=4 experiments). All data are represented as mean and standard error.

### Supplementary Tables

**Supplementary Table 1:** Demographic and clinical characteristics (mean ± SD) of the OOA study population.

| | **WES and Chip** | **WGS** |
|---|---|---|
| **N** | 5890 | 1083 |
| **Male (%)** | 43 | 50 |
| **Age (years)** | 41.8 ± 15.2 | 50.4 ± 16.8 |
| **BMI (kg/m$^2$)** | 26.6 ± 4.9 | 26.9 ± 4.5 |
| **LDL-C (mg/dl)** | 132.7 ± 45.8 | 140.4 ± 43.2 |
| **HDL-C (mg/dl)** | 61.4 ± 16.6 | 55.9 ± 15.6 |
| **TG (mg/dl)** | 71.5 ± 44.6 | 77.7 ± 48.8 |
| **Lipid-lowering Medication (%)** | 1.9 | 3.2 |

WES: Whole-exome sequencing
WGS: Whole-genome sequencing
BMI: Body Mass Index
LDL-C: Low density lipoprotein cholesterol
HDL-C: High density lipoprotein cholesterol
TG: triglyceride

79% of WGS samples were included in the WES and Chip samples

**Supplementary Table 2:** Top associated variants with LDL-C from whole exome sequencing.

| Locus | rsID | Chr:Position | Ref | Alt | MAF_Amish | Effect size | SE | P value | Gene | Known Locus |
|---|---|---|---|---|---|---|---|---|---|---|
| 1p13.3 | rs4970834 | 1:109272258 | C | T | 0.276 | -4.89 | 0.89 | 4.68E-08 | CELSR2 | Y |
| 6q25.3 | rs2282137 | 6:160047148 | G | A | 0.015 | 18.43 | 2.89 | 2.09E-10 | IGF2R | Y |
| 7p13 | rs217373 | 7:44572855 | C | A | 0.484 | -4.57 | 0.78 | 4.50E-09 | DDX56 | Y |
| 9p21.1 | rs551564683 | 9:33113783 | T | C | 0.060 | -13.49 | 1.69 | 1.65E-15 | B4GALT1 | N |
| 11q23.3 | rs11558589 | 11:119079170 | G | A | 0.029 | -14.92 | 2.26 | 4.23E-11 | VPS11 | Y |
| 17p11.2 | rs145316050 | 17:17819081 | G | A | 0.033 | -12.03 | 2.17 | 3.14E-08 | SREBF1 | Y |
| 19p13.2 | rs2228671 | 19:11100236 | C | T | 0.106 | -8.07 | 1.36 | 3.15E-09 | LDLR | Y |
| 19p13.11 | rs2304130 | 19:19678719 | A | G | 0.117 | -8.38 | 1.19 | 1.89E-12 | ZNF101 | Y |
| 19q13.32 | rs429358 | 19:44908684 | T | C | 0.136 | 7.42 | 1.08 | 6.93E-12 | APOE | Y |

Chr:Position: Chromosome:Position according to hg38
Ref: reference allele
Alt: alternative allele
MAF_Amish: minor allele frequency in the Amish
SE: Standard Error
Known Locus: previously reported in large lipid GWAS [29,43,47]

**Supplementary Table 3:** Top associated variants with LDL-C at the *B4GALT1* locus from whole genome sequencing.

| rsID | Position | Ref | Alt | $r^2$ | LDL_p-value_WGS | LDL_p-value_Imput | Fib_p-value_Imput | Info | Gene | Context | MAF_Amish | MAF_1KG_Eur |
|---|---|---|---|---|---|---|---|---|---|---|---|---|
| rs577483462 | 31523878 | A | G | 0.93 | 7.02E-06 | 4.25E-12 | 9.47E-04 | 0.994 | | intergenic | 0.059 | 0 |
| rs539837022 | 31599163 | G | A | 0.94 | 6.30E-06 | 5.81E-12 | 9.47E-04 | 0.995 | | intergenic | 0.059 | 0 |
| rs115335790 | 31853655 | G | A | 0.94 | 1.91E-05 | 6.53E-11 | 7.69E-04 | 0.994 | | intergenic | 0.059 | 0 |
| rs544554676 | 32231859 | C | T | 0.95 | 7.95E-06 | 8.10E-12 | 9.42E-04 | 0.995 | | intergenic | 0.059 | 0 |
| **rs551564683** | **33113783** | **T** | **C** | **---** | **3.14E-06** | **3.60E-15** | **8.44E-05** | **0.994** | **B4GALT1** | **missense** | **0.061** | **0** |
| rs1043957248 | 33892789 | G | C | 0.93 | 9.82E-06 | 1.32E-12 | 1.56E-04 | 0.991 | UBE2R2 | intron | 0.059 | 0 |
| rs1012682673 | 33971382 | G | A | 0.93 | 9.82E-06 | 1.87E-12 | 1.59E-04 | 0.991 | UBAP2 | intron | 0.059 | |
| rs998550643 | 33991396 | A | G | 0.93 | 9.82E-06 | 1.87E-12 | 1.59E-04 | 0.991 | UBAP2 | intron | 0.059 | |
| rs1023990285 | 34068625 | A | G | 0.93 | 9.82E-06 | 1.45E-12 | 2.58E-04 | 0.956 | | intergenic | 0.059 | |
| rs988733439 | 34277760 | CTA | C | 0.93 | 9.82E-06 | 7.48E-13 | 1.41E-04 | 0.982 | KIF24 | intron | 0.059 | 0 |
| rs898800582 | 34520553 | G | A | 0.93 | 9.82E-06 | 9.41E-13 | 1.51E-04 | 0.996 | DNAI1 | intron | 0.059 | |
| rs1040328811 | 34860997 | A | C | 0.92 | 2.35E-05 | 1.10E-11 | 1.21E-04 | 0.996 | | intergenic | 0.059 | |
| rs536902364 | 35178759 | A | G | 0.92 | 2.35E-05 | 4.36E-11 | 4.21E-04 | 0.996 | UNC13B | intron | 0.059 | 0.001 |
| rs1054917306 | 35570098 | A | T | 0.92 | 2.32E-05 | 6.83E-11 | 6.94E-04 | 0.997 | | regulatory region | 0.058 | |
| rs186441167 | 35868612 | G | A | 0.85 | 1.50E-05 | 1.90E-09 | 5.80E-03 | 1.000 | OR13J1 | downstream | 0.052 | 0.0089 |
| rs370946679 | 35816052 | A | G | 0.85 | 1.50E-05 | 1.90E-09 | 5.80E-03 | 1.000 | SPAG8 | upstream | 0.052 | 0.0179 |
| rs375489986 | 35816054 | T | C | 0.85 | 1.50E-05 | 1.90E-09 | 5.80E-03 | 1.000 | SPAG8 | upstream | 0.052 | 0.0179 |
| rs118137738 | 35857852 | G | A | 0.85 | 1.50E-05 | 1.90E-09 | 5.80E-03 | 1.000 | TMEM8B | downstream | 0.052 | 0.0169 |
| rs139585733 | 35875106 | G | A | 0.85 | 1.50E-05 | 1.90E-09 | 5.80E-03 | 1.000 | NDUFA5P4 | non_coding_transcript_exon | 0.052 | 0.0169 |
| rs569363002 | 35880024 | CA | C | 0.85 | 1.50E-05 | 1.91E-09 | 5.80E-03 | 1.000 | NDUFA5P4 | upstream | 0.052 | 0 |
| rs149557496 | 35906658 | G | A | 0.84 | 1.59E-05 | 1.61E-09 | 5.73E-03 | 0.999 | HRCT1 | 3 prime UTR | 0.052 | 0.0129 |

Position: variant position according to hg38
Ref: reference allele
Alt: alternative allele
$r^2$: linkage disequilibrium with the top associated variant rs551564683
LDL_p-value_WGS: LDL-C association analysis p-value from WGS (n=1,083)
LDL_p-value_Imput: LDL-C association analysis p-value from imputed data (n=5,890)
Fib_p-value_Imput: Fibrinogen association analysis p-value from imputed data (n=805)
Info: Imputation quality score
MAF_Amish: minor allele frequency in the Amish
MAF_1KG_Eur: minor allele frequency 1000 Genomes Project samples of European ancestry

**Supplementary Table 4:** The association between *B4GALT1* p.Asn352Ser and relevant traits in the OOA.

| TRAIT | N (total) | N (352Asn) | N (heterozygotes) | N (352Ser) | Effect size | p value |
|---|---|---|---|---|---|---|
| *Lipids* | | | | | | |
| Low-density Lipoprotein Cholesterol (mg/dL) | 5890 | 5194 | 680 | 16 | -13.49 | 1.65E-15 |
| High-density Lipoprotein Cholesterol (mg/dL) | 5890 | 5194 | 680 | 16 | -2.17 | 4.17E-03 |
| Triglycerides (mg/dL) | 5890 | 5194 | 680 | 16 | -2.57 | 1.32E-01 |
| *Liver enzymes* | | | | | | |
| Aspartate Transaminase (U/L) | 5595 | 4923 | 657 | 15 | 1.87 | 2.32E-11 |
| Alanine Aminotransferase (U/L) | 5795 | 5098 | 680 | 17 | 0.03 | 9.21E-01 |
| Alkaline Phosphatase (U/L) | 5795 | 5098 | 680 | 17 | 0.44 | 4.24E-01 |
| *Inflamatory markers* | | | | | | |
| C-Reactive Protein (mg/L) | 1338 | 1194 | 141 | 3 | 0.21 | 6.77E-01 |
| Interleukin 1 beta (pg/ml) | 711 | 645 | 65 | 1 | -0.13 | 8.22E-01 |
| Interleukin 6 (pg/ml) | 711 | 645 | 65 | 1 | 0.66 | 3.56E-01 |
| Matrix metallopeptidase 1 (ng/ml) | 712 | 646 | 65 | 1 | 0.40 | 5.78E-02 |
| Matrix Metallopeptidase 9 (ng/ml) | 712 | 646 | 65 | 1 | 58.04 | 6.01E-01 |
| White Blood Cell Count (Thousand/ul) | 5778 | 5086 | 675 | 17 | -0.01 | 8.74E-01 |
| *Calcification* | | | | | | |
| Coronary Calcification Score | 758 | 649 | 108 | 1 | 23.84 | 5.12E-01 |
| Aortic Calcification Score | 578 | 495 | 82 | 1 | -103.66 | 7.36E-01 |
| *Others* | | | | | | |
| Common Carotid Intima-Media Thickness (mm) | 1146 | 1024 | 119 | 3 | 3.98E-04 | 9.74E-01 |
| Liver to spleen density | 617 | 529 | 87 | 1 | 2.30E-02 | 3.19E-01 |
| Fibrinogen (mg/dL) | 805 | 721 | 81 | 3 | -26.62 | 9.87E-05 |

**Supplementary Table 5:** The ranges of GGT and coagulation measures in 12 subjects with *B4GALT1* 352Ser homozygotes.

| TRAIT | Subject range | Reference range | Notes |
|---|---|---|---|
| Gamma-Glutamyl Transpeptidase (U/L) | 8 - 27 | 3 - 65 | All normal |
| *Coagulation measures* | | | |
| activated Partial Thromboplastin Time (sec) | 27 - 35 | 22 - 34 | Two persons borderline high |
| Prothrombin Time (sec) | 10 - 12 | 9 - 11.5 | One person borderline high |
| Internationalized Normalized Ratio | 1 - 1.2 | 0.9 - 1.1 | One person borderline high |

**Supplementary Table 6:** Carbohydrate deficient transferrin test results from serum of 352Asn homozygotes (352Asn), heterozygotes (Asn352Ser) and 352Ser homozygotes (352Ser). (9 352Ser homozygotes, 8 Asn352Ser heterozygotes and 11 352Asn homozygotes)

| | Subject range | | | | Reference range | | |
|---|---|---|---|---|---|---|---|
| **Ratio** | **352Asn** | **Asn352Ser** | **352Ser** | **p-value** | **Normal** | **Indeterminate** | **Abnormal** |
| **Transferrin Mono-oligo/Di-oligo Ratio** | 0.03 - 0.05 | 0.04 - 0.08 | 0.03 - 0.06 | 0.0765 | < = 0.06 | 0.07-0.09 | > = 0.10 |
| **Transferrin A-oligo/Di-oligo Ratio** | 0.003 - 0.008 | 0.002 - 0.008 | 0.003 - 0.009 | 0.0597 | < = 0.011 | 0.012-0.021 | > = 0.022 |
| **Transferrin Tri-sialo/Di-oligo Ratio** | 0.03 - 0.06 | 0.04 - 0.13 | 0.17 - 0.3 | 9.12E-10 | < = 0.05 | 0.06-0.12 | > = 0.13 |
| **Apo CIII-1/Apo CIII-2 Ratio** | 1.05 - 2.63 | 1.15 - 2.36 | 0.95 - 2.33 | 0.358 | < = 2.91 | 2.92-3.68 | > = 3.69 |
| **Apo CIII-0/Apo CIII-2 Ratio** | 0.16 - 0.38 | 0.18 - 0.4 | 0.14 - 0.42 | 0.136 | < = 0.48 | 0.49-0.68 | > = 0.69 |

**Supplementary Table 7:** Mean (SD) of % peak area of significantly different glycans in plasma N-linked glycoproteins of *B4GALT1* 352Asn homozygotes (352Asn) and 352Ser homozygotes (352Ser) (n=12 per genotype group).

| Glycan | 352Asn | 352Ser | p-value |
|---|---|---|---|
| **G0F** | 0.34 (0.12) | 1.12 (0.2) | 5.42E-11 |
| **bG0** | 0.14 (0.11) | 1.21 (0.57) | 2.16E-06 |
| **G1S1** | 0.59 (0.21) | 3.86 (0.5) | 5.12E-16 |
| **G2S2** | 39.39 (1.68) | 32.88 (2.09) | 2.61E-08 |

**Supplementary Table 8:** Mean (SD) of % peak area of significantly different N-linked glycans in plasma ApoB100 of *B4GALT1* 352Asn homozygotes (352Asn) and 352Ser homozygotes (352Ser) (n=12 per genotype group).

| Glycan | 352Asn | 352Ser | p-value |
|---|---|---|---|
| **G0** | 0.0 (0.0) | 2.27 (0.44) | 1.13E-14 |
| **G1** | 0.0 (0.0) | 4.42 (0.81) | 4.47E-15 |
| **G1S1** | 1.47 (0.56) | 12.69 (0.90) | 3.21E-21 |
| **G2S1** | 29.24 (1.73) | 23.70 (1.10) | 3.96E-09 |
| **G2S2** | 32.70 (1.94) | 21.22 (1.29) | 3.56E-14 |

**Supplementary Table 9:** Mean (SD) of % peak area of significantly different N-linked glycans in plasma fibrinogen of *B4GALT1* 352Asn homozygotes (352Asn) and 352Ser homozygotes (352Ser) (n=12 per genotype group).

| Glycan | 352Asn | 352Ser | p-value |
|---|---|---|---|
| **G0** | 0.0 (0.0) | 1.15 (0.68) | 6.64E-06 |
| **G0F** | 1.11 ( 0.46) | 2.99 (0.63) | 3.19E-08 |
| **G1S1** | 2.37 (1.24) | 20.82 (6.64) | 3.31E-09 |
| **G2S1** | 47.44 (3.22) | 34.28 (4.25) | 1.94E-08 |
| **G2S2** | 28.06 (3.47) | 17.73 (4.73) | 3.92E-06 |

**Supplementary Table 10:** Mean (SD) of % peak area of significantly different N-linked glycans in plasma IgG of *B4GALT1* 352Asn homozygotes (352Asn) and 352Ser homozygotes (352Ser) (n=12 per genotype group).

| Glycan | 352Asn | 352Ser | p-value |
|---|---|---|---|
| **G0F** | 18.21 (5.37) | 38.90 (5.18) | 2.45E-09 |
| **G1F** | 26.59 ( 2.51) | 21.08 (3.48) | 2.05E-04 |
| **G2F** | 13.05 (3.7) | 2.78 (1.01) | 4.57E-09 |
| **G2FS1** | 10.97 (3.27) | 3.87 (0.98) | 3.21E-07 |
