## Supplemental Materials and Methods for "Genetic and functional evidence relates a missense variant in *B4GALT1* to lower LDL-C and fibrinogen"

[4]Full list of names and affiliations in the online supplementary materials

[5]Geriatrics Research and Education Clinical Center, Baltimore VA Medical Center, Baltimore, MD 21201, USA

### TABLE OF CONTENTS

#### Methods


| | |
|---|---|
| Study population and phenotypes | 3 |
| Whole-exome sequencing | 4 |
| Whole-genome sequencing | 4 |
| Chip genotyping and imputation | 4 |
| Association analyses | 5 |
| Carbohydrate deficient transferrin (CDT) test | 5 |
| Extraction of glycoproteins from human plasma | 5 |
| (HILIC-FLR-MS) glycoprotein analysis | 6 |
| Expression Constructs | 8 |
| Cell Lines and Media | 8 |
| Immunoprecipitation and glycosyltransferase activity assays | 8 |
| Measuring the enzymatic activities using HILIC-MS | 9 |
| Zebrafish *b4galt1* knock-down and phenotyping | 9 |

| | |
|---|---|
| **Data availability** | 10 |
| **References** | 11 |
| **Supplementary Materials** | |
| TOPMed Banner Authorship | 13 |
| Regeneron Genetics Center Banner Authorship | 22 |

### METHODS

#### Study population and phenotypes

The Old Order Amish (OOA) population of Lancaster County, PA immigrated to the Colonies from Central Europe in the early 1700's. There are currently around 40,000 OOA individuals in the Lancaster area, nearly all of whom can trace their ancestry back about 15 generations to approximately 750 founders. Investigators at the University of Maryland Baltimore have been studying the genetic determinants of cardiometabolic health in this population since 1993. To date, over 7,000 Amish adults have participated in one or more of our studies as part of the Amish Complex Disease Research Program[48], including the Heredity and Phenotype Intervention (HAPI) Heart Study[49], the Amish Longevity Study (ALS)[50], and the Amish Wellness Study[48]. These studies collected large numbers of variables including demographic and anthropometric information, medical history, clinical characteristics, lifestyle factors, and study specific variables, as well as blood and urine samples. All study protocols were approved by the institutional review board at the University of Maryland Baltimore. Informed consent was obtained from each study participant.

Fasting blood samples were used to measure lipids (total cholesterol, HDL-C, triglycerides), liver enzymes (ALT, AST, ALP, GGT), coagulation markers (fibrinogen, aPTT, PT, and INR), and inflammatory markers (CRP, WBC) (Quest Diagnostics, Horsham, PA). Other inflammatory markers were measured at the Cytokine Core Laboratory at the University of Maryland, Baltimore (IL1b, IL6, MMP1, MMP9). LDL-C was calculated using the Friedewald formula. For subjects on lipid lowering medication (~2% of research participants), LDL-C level was adjusted by dividing by 0.7.

The electron beam computerized tomography (EBCT) scans were performed on an Imatron C-150 scanner (GE, South San Francisco, Calif) in Timonium (Md). CAC scanning was performed using a standard protocol that included 30 to 40 three-mm contiguous transverse slices between the aortic root and the apex of the heart, gated to 80% of the RR interval and obtained during a single breathhold. The extent of calcium in the thoracic aorta was assessed by scanning between the superior aspect of the aortic arch and the superior pole of the kidney at 6-mm intervals. We elected to scan only the thoracic and upper abdominal aorta to limit radiation exposure. CAC was quantified using the Agatston method, which incorporates both density and area[51].The presence of calcification was defined as a density >130 Hounsfield units in >3 contiguous pixels (>1 mm2). The sum of the scores in the left main, left anterior descending, circumflex, and right coronary arteries was considered the CAC score. AC also was measured using the Agatston method, and the sum of all the AC lesions was considered the AC score. All scans were scored by a single experienced cardiologist using AccuImage (AccuImage Diagnostic Corp, San Francisco, Calif) software. Interscan reproducibility for quantification of CAC with this software was previously reported to range from 89%[52] to 94%[53]. The interreader and intrareader reproducibilities were each ≈99%[52]. Reproducibility of AccuImage measures of AC has not been reported; however, the median reproducibility of AC Agatston score using a similar scoring software system[54] was 90% with interreader and intrareader reproducibilities of 99% and 93%, respectively[54]. We defined presence of calcification as a CAC (or AC) score ≥1.

Measurements from two regions-of-interest, the liver and spleen, were also obtained. The spleen measurements were used as an attenuation standard. Accu View (Accuimage Corp.) software was used to calculate the attenuation coefficient in Hounsfield Units for each region-of-interest. Two 1.0-cm2 region-of-interest measurements were obtained from the liver and one was obtained from the spleen[55].

For carotid measures, B-mode ultrasonography was performed using a high-resolution linear transducer. Three frozen images of the left and right common carotid arteries were obtained concurrent to the R wave peaks on a simultaneous ECG recording. We analyzed a 10-mm segment of the far wall of the distal common carotid artery bilaterally using an automated edge detection system, which traces the lumen-intima interface and the media-adventia interface, which is then edited by the reader. The average distance between these interfaces serves as the IMT of the far wall. The average of the 2 measurements of the left common carotid artery IMT and 2 measurements for the right common carotid IMT (cIMT), obtained from 4 images, was used as a subject's mean common cIMT.

**Whole-exome sequencing**

Exome capture and sequencing were performed on 6,229 OOA samples at the Regeneron Genetics Center (RGC). Exome capture was performed using xGen design available from IDT with some modifications. The captured libraries were sequenced on the Illumina HiSeq 2500 platform with v4 chemistry using 75 bp paired-end reads. Paired-end sequencing of the captured bases was performed so that >85% of the bases were covered at 20x depth or greater, which is sufficient for calling heterozygous variants across most of the targeted bases. Read alignment and variant calling were performed using BWA-MEM and GATK as implemented in the RGC DNAseq analysis pipeline. SNPs with call rate <90%, and monomorphic SNPs were excluded. SNPs on X and Y chromosomes and the mitochondrial genome were also excluded. Samples failing QC metrics for contamination (N=8), low coverage (N=15), high levels of Mendelian errors (N=23), identical or MZ twins (one of each pair, N=23), withdrawn consent (N=5), gender mismatch (N=12), duplicates (N=12), or missing LDL-C or covariates (N=241) were excluded leaving 5,890 samples for the association analysis.

**Whole-genome sequencing**

Library preparation and whole-genome sequencing were performed on 1,113 OOA samples by the Broad Institute of MIT and Harvard. The NHLBI Informatics Resource Core at the University of Michigan performed alignment, base calling, and sequence quality scoring and variant calling of all TOPMed samples using the GotCloud pipeline[56]. Variant calling used a support vector machine (SVM) trained using known variants. Variants passing all quality filters with read depth at least 10 were delivered in BCF format and used for association analysis. Further variant QC included removing all sites in low-complexity regions [57], and on the X chromosome. We also removed variants with >5% missing rate, deviation from Hardy-Weinberg expectation (HWE) at p<1.0E-09, and variants with minor allele frequency (MAF) <0.1%. Sample QC was performed to remove samples with high levels of Mendelian errors (N=2), or identical or MZ twins (one of each pair) (N=4). Samples with missing LDL-C or covariates were also excluded (N=24) leaving 1,083 samples for the association analysis.

**Chip genotyping and imputation**

Genomic DNA was extracted from whole blood of 6,229 OOA and quantitated using picogreen. The chip genotyping for Infinium Global Screening Array v1.0 from Illumina that included 635,009 variants was performed at the Regeneron Genetic Center. Illumina iScan Infinium HTS workflow was used for analysis and genotype calling[58]. Prior to imputation, variants with >2% missing data (N=58,293), HWE < 1E-10 (N=571), Mendelian errors >1% (N=670) or with MAF < 0.01 (N=211,646) were excluded. We also excluded variants in the Y chromosome or mitochondrial genome (N=1,573), palindromic variants with frequency > 0.4 (N=344), and variants that were not in TOPMed freeze 5b reference panel (N=10,763). Sample QC was performed to remove samples with low quality (N=2), gender discordant (N=12), duplicate (N=10), high levels of Mendelian errors (N=25), more than 5% missing data (N=4), identical or MZ twins (one of each pair) (N=23). These QC procedures left 6,153 participants and 351,149 variants in the genotype file ready for imputation. The genotype data were uploaded to the Michigan Imputation Server where the pre-phasing was performed using Eagle v2.4, and then imputation to the TOPMed Freeze 5b reference panel was performed using Minimac4[59]. Following imputation, we excluded variants with imputation quality/INFO < 0.3, MAF <0.0001%or deviation from Hardy-Weinberg expectation at p< 1.0E-09.

Samples with missing LDL-C or covariates were excluded (N=254) leaving 5,890 samples for the association analysis.

**Association analyses**

Genetic association analyses were performed using linear mixed models to account for familial correlation using genetic relationship matrix (GRM)[60]. Analyses of LDL-C were adjusted for age, age squared, sex, Amish sub-study, and *APOB* p.Arg3527Gln. *APOB* p.Arg3527Gln is enriched in the Amish at 12% carrier frequency and increases LDL-C ~60 mg/dl [61], thus can confound LDL-C analysis if not included. The association analyses for fibrinogen, alkaline phosphatase, and natural log transformed AST and ALT were performed similarly adjusting for age, age squared, sex, Amish sub-study, and GRM. A p-value of 5.0E-08 was used to declare genome-wide significance for LDL-C. Bonferroni corrected p-value of 0.001 was used to correct for the 15 traits tested only for p.Asn352Ser, and testing fibrinogen for the other 20 correlated variants.

**Carbohydrate deficient transferrin (CDT) test**

The CDT test was performed using 0.1 ml serum samples from 28 participants from the 3 *B4GALT1* genotype groups (9 352Ser homozygotes, 8 Asn352Ser heterozygotes and 11 352Asn homozygotes). Each 352Ser homozygote was matched with a heterozygote and a 352Asn homozygote that were either siblings or closely related same sex individuals based on the kinship coefficient. They were also matched for age (within 5 years), and carrier status for *APOB* p.Arg3527Gln genotype. We were unable to enroll matches for one 352Ser homozygote, and 3 352Asn homozygotes. None of the participants were taking lipid-lowering medication.

Water-diluted samples were double washed using an immunoaffinity column. Glycosylation profiling of eluted proteins was performed using a mass spectrometer operated with 2 scan ranges specific for APOCIII and transferrin. Glycoform ratios of each protein were used to determine glycosylation deficiency. Glycoform ratios of transferrin include transferrin mono-oligo/Di-oligo ratio, transferrin A-oligo/Di-oligo ratio, and transferrin Tri-sialo/Di-oligo ratio. Glycoform ratios of APOCIII include ApoCIII-1/ApoCIII-2 ratio, and ApoCIII-0/ApoCIII-2 ratio

and apolipoprotein CIII (ApoCIII). The CDT test was performed at the Mayo medical laboratory of Mayo Clinic.

Mixed model analysis adjusted for age, sex, and GRM was used to test the association with the 5 measured ratios, and a p-value of 0.01 was considered significant.

**Extraction of glycoproteins from human plasma**

Glycoprotein analyses were performed using plasma samples from 24 participants (12 352Ser homozygotes and 12 352Asn homozygotes). Each 352Ser homozygote was matched with a 352Asn homozygote that was either a sibling or closely related same sex individual based on the kinship coefficient. They were also matched for age, and carrier status for *APOB* p.Arg3527Gln genotype. None of the participants were taking lipid-lowering medication. The difference between the 352Asn and 352Ser homozygotes for each glycan was assessed using a two-sided student t-test.

For isolation of global glycoproteins, aliquots of 50 µL of each human plasma sample were diluted 10 times with wash/bind buffer (0.15M NaCl, 1mM $MnCl_2$, 1mM $CaCl_2$, 20mM Tris, pH7.4) containing protease inhibitor cocktail. Diluted plasma samples were placed in a spin column containing equilibrated lectin resin (Agarose-bound Wheat Germ Agglutinin and Concanavalin A). Following standing at room temperature for 10 minutes, the mixture was centrifuged, and the retained resin was washed with wash/bind buffer (500 µL ×4), and then N-linked glycoprotein was eluted (500 µL ×4) with elution buffer (0.25 M Methyl-$\alpha$-D-mannopyranoside and 0.5 M N-acetyl-glucosamine in 20 mM Tris and 0.5 M NaCl, pH 7.4). The eluent was concentrated using a 3K MW cutoff spin filter, and resuspended in 40×2 µL of 50 mM HEPES buffer, pH 7.9 for N-linked glycan analysis using HILIC-FLR-MS.

For isolation of fibrinogen, aliquots of 25 µL of each human plasma sample were diluted 4 times with PBS containing protease inhibitor cocktail. Diluted plasma samples were placed in a spin column containing equilibrated Capture Select resin (Fibrinogen Affinity Matrix from Thermo Fisher Scientific) and 200 µL of PBS. Following incubating at room temperature in a rotator for 1 hour, the mixture was centrifuged, the retained resin was washed with PBS (200 µL ×4), and fibrinogen was eluted (200 µL ×2) with elution buffer (20 mM Tris, 50% [V/V] propylene glycol, 1 M arginine, pH 7.4) after incubated at room temperature for 10 minutes. The eluent was concentrated using a 3K MW cutoff spin filter, and resuspended in 40 µL of PBS for N-linked glycan analysis using HILIC-FLR-MS.

For isolation of total IgGs, aliquots of 15 µL of each human plasma sample were diluted 13 times with PBS containing protease inhibitor cocktail. Diluted plasma samples were placed in a spin column containing equilibrated rPotein A Sepharose Fast Flow slurry (GE Healthcare). Following incubating at room temperature for 5 minutes, the flow-through were collected by gravity. The retained resin was washed with PBS (300 µL ×3) by gently centrifuging, and then IgGs were eluted (200 µL ×3) with elution buffer (2.5% [V/V] acetic acid) by gently centrifuging. The eluent was concentrated using a 30K MW cutoff spin filter, and resuspended in 50 µL of PBS for N-linked glycan analysis using HILIC-FLR-MS.

To isolate ApoB100 by immunoprecipitation, aliquots of 25 µL of each human plasma sample were diluted by adding 25 µL of PBS (pH7.4) containing protease inhibitor cocktail. Diluted samples were placed in a 1.5 mL tube containing anti-ApoB100 antibody immobilized on magnetic Dynabeads (Thermo Fisher Scientific) equilibrated with BSA wash/binding buffer (0.1% BSA, 2 mM EDTA in PBS, pH 7.4). Following incubating at 4°C in a rotator for 1.5 hours, the mixture was placed on magnetic rack for 2 minutes, and the supernatant was collected in a new tube. The retained magnetic beads were washed with BSA wash/binding buffer (1 mL ×2) and PBS (1 mL ×2), and then ApoB100 was eluted with 50 µL of elution buffer (2% Waters RapiGest in PBS, pH 7.4) by incubating the mixture at 37°C for 20 minutes with shaking, then placing on the magnetic rack for 1 minute and removing the supernatant to a new tube for N-linked glycan analysis using HILIC-FLR-MS.

**Hydrophilic interaction liquid chromatography with fluorescence detection and mass spectrometry (HILIC-FLR-MS) analysis of global glycoproteins, fibrinogen, ApoB100 and IgG**

Aliquots (7.8 µL) of each eluate (global glycoproteins or fibrinogen) were transferred to a tube containing 15 µL DPBS with 2% RapiGest (Waters) and 1.2 µL of 0.1 M TCEP, and incubated at 80ºC for 10 minutes. After cooling to room temperature, 6 µL of peptide N-glycosidase F (PNGase F stock, 65 NEB unit/IUB milliunit/µL) was added and the mixture was incubated at 50ºC for 5 minutes to release N-linked glycans. Samples were cooled to room temperature and subjected to spin down condensation. Released glycans were fluorescently labeled by the addition of 12 µL of derivatization solution (9 mg *Rapi*Fluor-MS™ dissolved in 130 µL anhydrous *N-N*-Dimethylformamide [DMF] from Waters), and incubated at 50°C for 30 minutes. The derivatized glycans were diluted by addition of DMF (final concentration of 25% [v/v] DMF), acetonitrile (final concentration of 53% [v/v] acetonitrile) and Milli-Q water.

Aliquots (10 µL) of each eluate of IgGs were transferred to a tube containing 2 µL of 0.5 M HEPES (pH 7.9) with 1% RapiGest (Waters) and 2 µL of 50 mM TCEP, and incubated at 80ºC for 10 minutes. After cooling to room temperature, 6 µL of peptide N-glycosidase F (PNGase F stock, 65 NEB unit/IUB milliunit/µL) was added and the mixture was incubated at 45ºC for 25 minutes to release N-linked glycans. Samples were cooled to room temperature and subjected to spin down condensation. Released glycans were fluorescently labeled by the addition of 8 µL of derivatization solution (9 mg RapiFluor-MS™ dissolved in 130 µL DMF from Waters), and incubated at 45°C for 30 minutes. The derivatized glycans were diluted by addition of DMF (final concentration of 25% [v/v] DMF), acetonitrile (final concentration of 53% [v/v] acetonitrile) and Milli-Q water.

Aliquots (13 µL) of each eluate of ApoB-100 were transferred to a tube containing 11.8 µL DPBS with 2% RapiGest (Waters) and 1.2 µL of 0.1 M TCEP, and incubated at 80ºC for 10 minutes. After cooling to room temperature, 3 µL Lys-C Stock (0.1 µg/µL in Milli-Q water) was added and incubate at 37°C for 30 minutes, then 6 µL of peptide N-glycosidase F (PNGase F stock, 65 NEB unit/IUB milliunit/µL) was added, and the mixture was incubated at 50ºC for 5 minutes to release N-linked glycans. Samples were cooled to room temperature and subjected to spin down condensation. Released glycans were fluorescently labeled by the addition of 12 µL of derivatization solution (9 mg *Rapi*Fluor-MS™ dissolved in 130 µL DMF from Waters), and incubated at 50°C for 5 minutes. After added 1 µL of formic acid (FA), the mixtures were incubated at 50°C for 25 minutes. The derivatized glycans were diluted by addition of DMF (final concentration of 25% [v/v] DMF), acetonitrile (final concentration of 53% [v/v] acetonitrile) and Milli-Q water.

For HILIC separation of the glycans from global glycoprotein, fibrinogen and ApoB-100, a Waters UPLC Glycan BEH® Amide column (130Å, 1.7 µm, 2.1 mm × 150 mm) was used with a column temperature of 60°C and a solvent flow rate of 0.45 mL/minute. Mobile phase A was 50 mM ammonium formate in water, pH 4.4, and mobile phase B was 100% acetonitrile. A 50 µL aliquot of each diluted samples was injected onto the UPLC Glycan BEH® Amide column for HILIC-FLR-MS analysis. The column was pre-equilibrated in 25% mobile phase A. After sample injection, the percentage of mobile phase A was increased from 25% to 32.2% mobile phase A over 29 minutes followed by an additional increase to 45% mobile phase A over an additional 36 minutes to optimize oligosaccharide separation. The *Rapi*Fluor-MS™ labeled glycans were detected using a fluorescence detector set at an excitation wavelength of 265 nm and emission wavelength of 425 nm. The Thermo Q Exactive Plus mass spectrometer was operated in positive mode with a spray voltage of 4.0 kV, and sheath and auxiliary gas at 40 and 10 (arbitrary units), respectively. The mass spectra were acquired with an m/z range between 650 and 2000. Each glycoform of released N-linked glycans was quantitated by relative quantification (percentage peak area [%PA]) based on the FLR chromatograms. The total levels of galactosylation and sialylation were measured as: $[\sum(\%PA^* X_1)/2] + [\sum(\%PA^*X_2)/3]$; Where %PA is percentage peak area; $X_1$ = 1 or 2 galactose or sialic acid ; $X_2$ = 1, 2, or 3 galactose or sialic acid; 2, 3 = number of branches on glycan. The total levels of fucosylation were measured as: percentage peak area sum of all fucosylated glycans. Two-sided student t-test was used to compare the two opposite homozygote groups.

**Expression Constructs**

cDNA encoding wild-type Myc-FLAG-tagged *B4GATL1* (352Asn) was excised from the RC-213879 plasmid using sequential enzyme reactions consisting FseI digestion, Klenow treatment to produce blunt ends, and EcoRI digestion. The resulting 1.5 kb fragment was directionally cloned into the SmaI-EcoRI digested pIRES2-EGFP plasmid. Site-directed mutagenesis to generate the cDNA encoding a Myc-FLAG-tagged *B4GATL1* 352Ser variant was carried out using the following forward and reverse primers (5′-3′):
P4291-g (TGAACCCA**G**TCCTCAGAGGTTTGAC)
P4291-c (CTGAGGA**C**TGGGTTCATTTTTCTTG)

**Cell Lines and Media**

COS-7 cells and Huh-7 cells were obtained from the American Type Culture Collection. Cell cultures were maintained in Dulbecco's modified Eagle's medium (Invitrogen) supplemented with 10% fetal bovine serum, antibiotics (50 units/ml penicillin and 50 µg/ml streptomycin), and 1.0 mM sodium pyruvate (#11360070, Invitrogen).

**Immunoprecipitation and glycosyltransferase activity assays**

COS-7 cells were seeded in 6-well plates at $1x10^6$ cells per well 24 hours prior to transfection. Cells were transiently transfected with 3 ug of plasmid containing cDNA encoding myc-FLAG tagged *B4GALT1*- 352Asn or -352Ser, using Lipofectamine 2000 (#11668019, ThermoFisher) according to the manufacture’s protocol. After 48 hours cells were washed with PBS and scraped into lysis buffer containing 50 mM Tris-HCl, pH 7.4, 150 mM NaCl, 1mM EDTA, 1% Triton X-100, and protease inhibitors (#11836170001, Sigma). After incubation for 30 min, on ice, cells were subjected to sonication and centrifugation. Cleared cellular lysates were incubated with 40 µL of pre-washed Anti-FLAG M2 affinity gel suspension (A2220, Sigma) for 2 hours at 4°C. After incubation, beads were isolated by centrifugation and washed with 1X TBS for a total of three times. Bound protein complexes were immediately assayed for glycosyltransferase activity using the glycosyltransferase activity kit (EA001, R&D). The reaction mixture consisted of 25 µL of 1X reaction buffer (25 mM Tris, 10 mM $CaCl_2$, pH 7.5 and 10 mM $MnCl_2$), 10 µL of 0.5mM UDP-Galactose donor (#670111, Millipore), 10 µL of 50 mM N-Acetyl-D-glucosamine acceptor (#1079, Millipore) and 5 µL Coupling Phosphatase (20 ng/ µL; #895404, R&D) added directly to the beads. After incubation for 1 hour at 37°C, beads were pelleted, and supernatants were transferred to 96-well plates for malachite green colorimetric assay. Optical density was measured at 620 nm using a Tecan Spark plate reader. All samples were assayed in duplicate. A standard curve for recombinant human B4GALT1 protein (3609-GT, R&D) was generated according to the protocol specified by the glycosyltransferase activity kit, and the specific activity of the enzyme was calculated. Proteins on the retained immunoprecipitation beads were recovered by addition of 20 µL sample buffer containing beta-mercaptoethanol and boiling. Recovered proteins were subjected to SDS-PAGE and western blot analysis using the anti-myc (#2278, Cell Signaling) antibody. The data are expressed as a ratio of B4GALT1 enzyme activity to the amount of 352Asn or 352Ser recovered and quantified by densitometry.

#### Measuring the enzymatic activities using hydrophilic interaction liquid chromatography with mass spectrometry (HILIC-MS) analysis

Recombinant human 352Asn and Ser352 B4GALT1 proteins were synthesized by R&D Systems. The proteins' enzymatic activities were measured using uridine diphosphate galactose (UDP-Gal) and *N*-acetylglucosamine (GlcNAc) as the substrates over eight different time points in triplicate. The assay (reaction) buffer contains 25 mM Tris pH7.5, 150 mM sodium chloride, 10 mM magnesium chloride, and 10 mM manganese chloride. The enzymatic reaction was initiated by adding 1.1 pmol of each B4GALT1 into the mixture of 1.7 nmol UDP-Gal, and 500 nmol GlcNAc, and the mixture was incubated at room temperature for 2, 5, 10, 15, 25, 40, and 60 minutes. The sample representing the initial time point (0 min) was prepared by omitting the enzyme. A HILIC-MS based method was used to quantitate the decrease of UDP-Gal. Samples were analyzed using an Acquity UPLC I-Class coupled with Vion IMS Q-Tof (Waters, Milford, MA, USA). A Waters Acquity UPLC Glycoprotein BEH Amide chromatographic column (300 Å, 150 mm × 2.1 mm, 1.7 µm) was used with a column temperature of 45°C and a solvent flow rate of 0.2 mL/minute. Mobile phase A was 50 mM ammonium acetate pH 7.5, and mobile phase B was 5% (v/v) 25 mM ammonium acetate pH 7.5 in acetonitrile. A 1 µL aliquot of each mixture was injected onto the column for HILIC-MS analysis. The column was pre-equilibrated in 15% mobile phase A. After sample injection, the percentage of mobile phase A was increased from 15% to 33% mobile phase A over 1 minute followed by an additional increase to 36% mobile phase A over an additional 10 minutes to optimize separation and UDP-Gal elution. The samples were monitored using a TUV detector at a wavelength of 280 nm. The Waters Vion IMS Q-Tof mass spectrometer was operated in positive mode and the mass spectra were acquired with a m/z range between 100 and 1000. UDP-Gal was quantitated by measurement of UV peak areas based on the UV chromatograms and the data were collected and processed using UNIFI (Waters).

#### Zebrafish *b4galt1* knock-down and phenotyping

Wild-type (Tubingen) zebrafish stocks were used to generate embryos for morpholino or mRNA injection. Adult fish were maintained and bred at 27-29°C and embryos were raised at 28.5°C. All animals were housed and maintained in accordance with protocols approved by the University of Maryland Baltimore Institutional Animal Care and Use Committee. Morpholino antisense oligonucleotides (MOs) were obtained (Gene Tools, Inc.) based on previously published MOs targeted against *b4galt1*[27] or *ldlr* MO[25]. 8ng or 10ng of b4galt1 MO1, 2ng or 4ng of b4galt1 MO2, or 5ng of *ldlr* MO were injected at the 1-2 cell stage and *b4galt1* MOs validated by qRT-PCR quantification of endogenous *b4galt1* transcript. Off-target toxicity was assessed by qRT-PCR quantification of the delta113 isoform of p53[62]. For mRNA rescue experiments, human full length capped *B4GALT1* mRNA was transcribed via *in vitro* transcription from a pCS2+ plasmid vector containing the open reading frame (ORF) of the 352Asn or 352Ser *B4GALT1*. mRNA was mixed with MO at varying concentrations and co-injected into 1-2 cell stage embryos. Between 25-150pg of 352Asn *B4GALT1* mRNA was co-injected with 8ng *b4galt1* MO1 for LDL-C quantification assays to identify optimal rescue concentration. 100pg of 352Asn or 352Ser *B4GALT1* mRNA were co-injected with 8ng *b4galt1* MO1 for LDL-C rescue experiments. For each injection experiment, a total of 200-400 embryos were injected and each experiment was repeated a minimum of three times.

For LDL-C quantification, we homogenized 100 5-day post-fertilization (dpf) larvae per experiment in 400 μl of ice-cold 10 μM butylated hydroxytoluene and filtered the homogenate through a 0.45-μm Dura PVDF membrane filter (Millipore) in preparation for lipid extraction. Using the HDL and LDL/VLDL Cholesterol Assay Kit (Cell Biolabs, Inc.) we processed the homogenate as per manufacturer's protocol. After precipitation and dilution, samples were analyzed by fluorimetric analysis using a SpectraMax Gemini EM plate reader and SoftMax Pro microplate data acquisition and analysis software (Molecular Devices). Two-sided student t-test was used for pairwise comparison between groups.

Fat content in larval livers was examined at 5 dpf, via Oil Red O staining of whole animals fixed in 4% paraformaldehyde (PFA). Hepatic lipid was assessed by quantification of stained hepatic lipid droplets and identifying proportions of larvae in each experimental having no (zero), low (<10 droplets), medium (<20), or high (>20) droplets.

### Data availability

The Old Order Amish TOPMed whole-genome sequencing, phenotype and covariate data used in this report are available through dbGaP. The study name is NHLBI TOPMed: Genetics of Cardiometabolic Health in the Amish and accession is phs000956.v3.p1. Whole-exome sequencing, chip genotypes, and imputed data are property of Regeneron Genetics Center,LLC. Glycoprotein profiling is property of Regeneron Pharmaceuticals, Inc. Targeted genotyping and CDT test results are available from the University of Maryland Baltimore investigators on request to qualified academic researchers. Zebrafish gRNA sequences, MO sequences and primer sequences for qRT-PCR available upon request.

**Supplementary Materials**

1- TOPMed Banner Authorship

| Name | Institution(s) |
|---|---|
| Abe, Namiko | New York Genome Center |
| Abecasis, Goncalo | University of Michigan |
| Albert, Christine | Massachusetts General Hospital |
| Allred, Nicholette (Nichole) Palmer | Wake Forest Baptist Health |
| Almasy, Laura | Children's Hospital of Philadelphia, University of Pennsylvania |
| Alonso, Alvaro | Emory University |
| Ament, Seth | University of Maryland |
| Anderson, Peter | |
| Anugu, Pramod | University of Mississippi |
| Applebaum-Bowden, Deborah | National Institutes of Health |
| Arking, Dan | Johns Hopkins University |
| Arnett, Donna K | University of Kentucky |
| Ashley-Koch, Allison | Duke University |
| Aslibekyan, Stella | University of Alabama |
| Assimes, Tim | Stanford University |
| Auer, Paul | University of Wisconsin Milwaukee |
| Avramopoulos, Dimitrios | Johns Hopkins University |
| Barnard, John | Cleveland Clinic |
| Barnes, Kathleen | University of Colorado at Denver |
| Barr, R. Graham | Columbia University |
| Barron-Casella, Emily | Johns Hopkins University |
| Beaty, Terri | Johns Hopkins University |
| Becker, Diane | Johns Hopkins University |
| Becker, Lewis | Johns Hopkins University |
| Beer, Rebecca | National Heart, Lung, and Blood Institute, National Institutes of Health |
| Begum, Ferdouse | Johns Hopkins University |
| Beitelshees, Amber | University of Maryland |
| Benjamin, Emelia | Boston University, Massachusetts General Hospital |
| Bezerra, Marcos | Fundação de Hematologia e Hemoterapia de Pernambuco - Hemope |
| Bielak, Larry | University of Michigan |
| Bis, Joshua | University of Washington |
| Blackwell, Thomas | University of Michigan |
| Blangero, John | University of Texas Rio Grande Valley School of Medicine |
| Boerwinkle, Eric | University of Texas Health at Houston |
| Borecki, Ingrid | University of Washington |

| Name | Institution(s) |
|---|---|
| Bowler, Russell | National Jewish Health |
| Brody, Jennifer | University of Washington |
| Broeckel, Ulrich | Medical College of Wisconsin |
| Broome, Jai | University of Washington |
| Bunting, Karen | New York Genome Center |
| Burchard, Esteban | University of California, San Francisco |
| Cardwell, Jonathan | University of Colorado at Denver |
| Carty, Cara | Women's Health Initiative |
| Casaburi, Richard | University of California, Los Angeles |
| Casella, James | Johns Hopkins University |
| Chaffin, Mark | The Broad Institute |
| Chang, Christy | University of Maryland |
| Chasman, Daniel | Brigham & Women's Hospital |
| Chavan, Sameer | University of Colorado at Denver |
| Chen, Bo-Juen | New York Genome Center |
| Chen, Wei-Min | University of Virginia |
| Chen, Yii-Der Ida | Los Angeles Biomedical Research Institute |
| Cho, Michael | Brigham & Women's Hospital |
| Choi, Seung Hoan | The Broad Institute |
| Chuang, Lee-Ming | National Taiwan University |
| Chung, Mina | Cleveland Clinic |
| Cornell, Elaine | University of Vermont |
| Correa, Adolfo | University of Mississippi |
| Crandall, Carolyn | University of California, Los Angeles |
| Crapo, James | National Jewish Health |
| Cupples, L Adrienne | Boston University |
| Curran, Joanne | University of Texas Rio Grande Valley School of Medicine |
| Curtis, Jeffrey | University of Michigan |
| Custer, Brian | Blood Systems Research Institute UCSF |
| Damcott, Coleen | University of Maryland |
| Darbar, Dawood | University of Illinois at Chicago |
| Das, Sayantan | University of Michigan |
| David, Sean | Stanford University |
| Davis, Colleen | |
| Daya, Michelle | University of Colorado at Denver |
| de Andrade, Mariza | Mayo Clinic |
| DeBaun, Michael | Vanderbilt University |
| Deka, Ranjan | University of Cincinnati |
| DeMeo, Dawn | Brigham & Women's Hospital |
| Devine, Scott | University of Maryland |

| Name | Institution(s) |
| --- | --- |
| Do, Ron | Icahn School of Medicine at Mount Sinai |
| Duan, Qing | University of North Carolina |
| Duggirala, Ravi | University of Texas Rio Grande Valley School of Medicine |
| Durda, Peter | University of Vermont |
| Dutcher, Susan | Washington University in St Louis |
| Eaton, Charles | Brown University |
| Ekunwe, Lynette | University of Mississippi |
| Ellinor, Patrick | Massachusetts General Hospital |
| Emery, Leslie | University of Washington |
| Farber, Charles | University of Virginia |
| Farnam, Leanna | Brigham & Women's Hospital |
| Fingerlin, Tasha | National Jewish Health |
| Flickinger, Matthew | University of Michigan |
| Fornage, Myriam | University of Texas Health at Houston |
| Franceschini, Nora | University of North Carolina |
| Fu, Mao | University of Maryland |
| Fullerton, Stephanie M. | University of Washington |
| Fulton, Lucinda | Washington University in St Louis |
| Gabriel, Stacey | The Broad Institute |
| Gan, Weiniu | National Heart, Lung, and Blood Institute, National Institutes of Health |
| Gao, Yan | University of Mississippi |
| Gass, Margery | Fred Hutchinson Cancer Research Center |
| Gelb, Bruce | Icahn School of Medicine at Mount Sinai |
| Geng, Xiaoqi (Priscilla) | University of Michigan |
| Germer, Soren | New York Genome Center |
| Gignoux, Chris | Stanford University |
| Gladwin, Mark | University of Pittsburgh |
| Glahn, David | Yale University |
| Gogarten, Stephanie | University of Washington |
| Gong, Da-Wei | University of Maryland |
| Goring, Harald | University of Texas Rio Grande Valley School of Medicine |
| Gu, C. Charles | Washington University in St Louis |
| Guan, Yue | University of Maryland |
| Guo, Xiuqing | Los Angeles Biomedical Research Institute |
| Haessler, Jeff | Fred Hutchinson Cancer Research Center, Women's Health Initiative |
| Hall, Michael | University of Mississippi |
| Harris, Daniel | University of Maryland |
| Hawley, Nicola | Yale University |

| Name | Institution(s) |
|---|---|
| He, Jiang | Tulane University |
| Heavner, Ben | University of Washington |
| Heckbert, Susan | University of Washington |
| Hernandez, Ryan | University of California, San Francisco |
| Herrington, David | Wake Forest Baptist Health |
| Hersh, Craig | Brigham & Women's Hospital |
| Hidalgo, Bertha | University of Alabama |
| Hixson, James | University of Texas Health at Houston |
| Hokanson, John | University of Colorado at Denver |
| Holly, Kramer | Loyola University |
| Hong, Elliott | University of Maryland |
| Hoth, Karin | University of Iowa |
| Hsiung, Chao (Agnes) | National Health Research Institute Taiwan |
| Huston, Haley | Blood Works Northwest |
| Hwu, Chii Min | Taichung Veterans General Hospital Taiwan |
| Irvin, Marguerite Ryan | University of Alabama |
| Jackson, Rebecca | Ohio State University Wexner Medical Center |
| Jain, Deepti | University of Washington |
| Jaquish, Cashell | National Heart, Lung, and Blood Institute, National Institutes of Health |
| Jhun, Min A | University of Michigan |
| Johnsen, Jill | Blood Works Northwest, University of Washington |
| Johnson, Andrew | National Heart, Lung, and Blood Institute, National Institutes of Health |
| Johnson, Craig | University of Washington |
| Johnston, Rich | Emory University |
| Jones, Kimberly | Johns Hopkins University |
| Kang, Hyun Min | University of Michigan |
| Kaplan, Robert | Albert Einstein College of Medicine |
| Kardia, Sharon | University of Michigan |
| Kathiresan, Sekar | The Broad Institute |
| Kaufman, Laura | Brigham & Women's Hospital |
| Kelly, Shannon | Blood Systems Research Institute UCSF |
| Kenny, Eimear | Icahn School of Medicine at Mount Sinai |
| Kessler, Michael | University of Maryland |
| Khan, Alyna | University of Washington |
| Kinney, Greg | University of Colorado at Denver |
| Konkle, Barbara | Blood Works Northwest |
| Kooperberg, Charles | Fred Hutchinson Cancer Research Center |
| Krauter, Stephanie | University of Washington |

| Name | Institution(s) |
|---|---|
| Lange, Christoph | Harvard School of Public Health |
| Lange, Ethan | University of Colorado at Denver |
| Lange, Leslie | University of Colorado at Denver |
| Laurie, Cathy | University of Washington |
| Laurie, Cecelia | University of Washington |
| LeBoff, Meryl | Brigham & Women's Hospital |
| Lee, Seunggeun Shawn | University of Michigan |
| Lee, Wen-Jane | Taichung Veterans General Hospital Taiwan |
| LeFaive, Jonathon | University of Michigan |
| Levine, David | University of Washington |
| Levy, Dan | National Heart, Lung, and Blood Institute, National Institutes of Health |
| Lewis, Joshua | University of Maryland |
| Li, Yun | University of North Carolina |
| Lin, Honghuang | Boston University |
| Lin, Keng Han | University of Michigan |
| Liu, Simin | Brown University, Women's Health Initiative |
| Liu, Yongmei | Wake Forest Baptist Health |
| Loos, Ruth | Icahn School of Medicine at Mount Sinai |
| Lubitz, Steven | Massachusetts General Hospital |
| Lunetta, Kathryn | Boston University |
| Luo, James | National Heart, Lung, and Blood Institute, National Institutes of Health |
| Mahaney, Michael | University of Texas Rio Grande Valley School of Medicine |
| Make, Barry | Johns Hopkins University |
| Manichaikul, Ani | University of Virginia |
| Manson, JoAnn | Brigham & Women's Hospital |
| Margolin, Lauren | The Broad Institute |
| Martin, Lisa | George Washington University |
| Mathai, Susan | University of Colorado at Denver |
| Mathias, Rasika | Johns Hopkins University |
| McArdle, Patrick | University of Maryland |
| McDonald, Merry-Lynn | University of Alabama |
| McFarland, Sean | Harvard University |
| McGarvey, Stephen | Brown University |
| Mei, Hao | University of Mississippi |
| Meyers, Deborah A | University of Arizona |
| Mikulla, Julie | National Heart, Lung, and Blood Institute, National Institutes of Health |
| Min, Nancy | University of Mississippi |

| Name | Institution(s) |
|---|---|
| Minear, Mollie | National Heart, Lung, and Blood Institute, National Institutes of Health |
| Minster, Ryan L | University of Pittsburgh |
| Mitchell, Braxton | University of Maryland |
| Montasser, May E. | University of Maryland |
| Musani, Solomon | University of Mississippi |
| Mwasongwe, Stanford | University of Mississippi |
| Mychaleckyj, Josyf C | University of Virginia |
| Nadkarni, Girish | Icahn School of Medicine at Mount Sinai |
| Naik, Rakhi | Johns Hopkins University |
| Natarajan, Pradeep | The Broad Institute, Harvard University, Massachusetts General Hospital |
| Nekhai, Sergei | Howard University |
| Nickerson, Deborah | University of Washington |
| North, Kari | University of North Carolina |
| O'Connell, Jeff | University of Maryland |
| O'Connor, Tim | University of Maryland |
| Ochs-Balcom, Heather | University at Buffalo |
| Pankow, James | University of Minnesota |
| Papanicolaou, George | National Heart, Lung, and Blood Institute, National Institutes of Health |
| Parker, Margaret | Brigham & Women's Hospital |
| Parsa, Afshin | University of Maryland |
| Penchev, Sara | National Jewish Health |
| Peralta, Juan Manuel | University of Texas Rio Grande Valley School of Medicine |
| Perez, Marco | Stanford University |
| Perry, James | University of Maryland |
| Peters, Ulrike | Fred Hutchinson Cancer Research Center, University of Washington |
| Peyser, Patricia | University of Michigan |
| Phillips, Larry | Emory University |
| Phillips, Sam | University of Washington |
| Pollin, Toni | University of Maryland |
| Post, Wendy | Johns Hopkins University |
| Powers Becker, Julia | University of Colorado at Denver |
| Preethi Boorgula, Meher | University of Colorado at Denver |
| Preuss, Michael | Icahn School of Medicine at Mount Sinai |
| Prokopenko, Dmitry | Harvard University |
| Psaty, Bruce | University of Washington |
| Qasba, Pankaj | National Heart, Lung, and Blood Institute, National Institutes of Health |

| Name | Institution(s) |
|---|---|
| Qiao, Dandi | Brigham & Women's Hospital |
| Qin, Zhaohui | Emory University |
| Rafaels, Nicholas | University of Colorado at Denver |
| Raffield, Laura | University of North Carolina |
| Ramachandran, Vasan | Boston University |
| Rao, D.C. | Washington University in St Louis |
| Rasmussen-Torvik, Laura | Northwestern University |
| Ratan, Aakrosh | University of Virginia |
| Redline, Susan | Brigham & Women's Hospital |
| Reed, Robert | University of Maryland |
| Regan, Elizabeth | National Jewish Health |
| Reiner, Alex | Fred Hutchinson Cancer Research Center, University of Washington |
| Rice, Ken | University of Washington |
| Rich, Stephen | University of Virginia |
| Roden, Dan | Vanderbilt University |
| Roselli, Carolina | The Broad Institute |
| Rotter, Jerome | Los Angeles Biomedical Research Institute |
| Ruczinski, Ingo | Johns Hopkins University |
| Russell, Pamela | University of Colorado at Denver |
| Ruuska, Sarah | Blood Works Northwest |
| Ryan, Kathleen | University of Maryland |
| Sakornsakolpat, Phuwanat | Brigham & Women's Hospital |
| Salimi, Shabnam | University of Maryland |
| Salzberg, Steven | Johns Hopkins University |
| Sandow, Kevin | Los Angeles Biomedical Research Institute |
| Sankaran, Vijay | Harvard University |
| Scheller, Christopher | University of Michigan |
| Schmidt, Ellen | University of Michigan |
| Schwander, Karen | Washington University in St Louis |
| Schwartz, David | University of Colorado at Denver |
| Sciurba, Frank | University of Pittsburgh |
| Seidman, Christine | Harvard Medical School |
| Sheehan, Vivien | Baylor College of Medicine |
| Shetty, Amol | University of Maryland |
| Shetty, Aniket | University of Colorado at Denver |
| Sheu, Wayne Hui-Heng | Taichung Veterans General Hospital Taiwan |
| Shoemaker, M. Benjamin | Vanderbilt University |
| Silver, Brian | UMass Memorial Medical Center |
| Silverman, Edwin | Brigham & Women's Hospital |

| Name | Institution(s) |
|---|---|
| Smith, Jennifer | University of Michigan |
| Smith, Josh | University of Washington |
| Smith, Nicholas | University of Washington |
| Smith, Tanja | New York Genome Center |
| Smoller, Sylvia | Albert Einstein College of Medicine |
| Snively, Beverly | Wake Forest Baptist Health |
| Sofer, Tamar | Brigham & Women's Hospital |
| Sotoodehnia, Nona | University of Washington |
| Stilp, Adrienne | University of Washington |
| Streeten, Elizabeth | University of Maryland |
| Sung, Yun Ju | Washington University in St Louis |
| Sylvia, Jody | Brigham & Women's Hospital |
| Szpiro, Adam | University of Washington |
| Sztalryd, Carole | University of Maryland |
| Taliun, Daniel | University of Michigan |
| Tang, Hua | Stanford University |
| Taub, Margaret | Johns Hopkins University |
| Taylor, Kent | Los Angeles Biomedical Research Institute |
| Taylor, Simeon | University of Maryland |
| Telen, Marilyn | Duke University |
| Thornton, Timothy A. | University of Washington |
| Tinker, Lesley | Women's Health Initiative |
| Tirschwell, David | University of Washington |
| Tiwari, Hemant | University of Alabama |
| Tracy, Russell | University of Vermont |
| Tsai, Michael | University of Minnesota |
| Vaidya, Dhananjay | Johns Hopkins University |
| VandeHaar, Peter | University of Michigan |
| Vrieze, Scott | University of Colorado at Boulder, University of Minnesota |
| Walker, Tarik | University of Colorado at Denver |
| Wallace, Robert | University of Iowa |
| Walts, Avram | University of Colorado at Denver |
| Wan, Emily | Brigham & Women's Hospital |
| Wang, Fei Fei | University of Washington |
| Watson, Karol | University of California, Los Angeles |
| Weeks, Daniel E. | University of Pittsburgh |
| Weir, Bruce | University of Washington |
| Weiss, Scott | Brigham & Women's Hospital |
| Weng, Lu-Chen | Massachusetts General Hospital |
| Willer, Cristen | University of Michigan |

| Name | Institution(s) |
|---|---|
| Williams, Kayleen | University of Washington |
| Williams, L. Keoki | Henry Ford Health System |
| Wilson, Carla | Brigham & Women's Hospital |
| Wilson, James | University of Mississippi |
| Wong, Quenna | University of Washington |
| Xu, Huichun | University of Maryland |
| Yanek, Lisa | Johns Hopkins University |
| Yang, Ivana | University of Colorado at Denver |
| Yang, Rongze | University of Maryland |
| Zaghloul, Norann | University of Maryland |
| Zekavat, Maryam | The Broad Institute |
| Zhang, Yingze | University of Pittsburgh |
| Zhao, Snow Xueyan | National Jewish Health |
| Zhao, Wei | University of Michigan |
| Zheng, Xiuwen | University of Washington |
| Zhi, Degui | University of Texas Health at Houston |
| Zhou, Xiang | University of Michigan |
| Zody, Michael | New York Genome Center |
| Zoellner, Sebastian | University of Michigan |

**2- Regeneron Genetics Center Banner Authorship**

All authors are listed in alphabetical order.

**RGC Management and Leadership Team**

Goncalo Abecasis, Ph.D., Aris Baras, M.D., Michael Cantor, M.D., Giovanni Coppola, M.D., Aris Economides, Ph.D., John D. Overton, Ph.D., Jeffrey Reid, Ph.D., Alan Shuldiner, M.D.

**Sequencing and Lab Operations**

Christina Beechert, Caitlin Forsythe, M.S., Erin D. Fuller, Zhenhua Gu, M.S., Michael Lattari, Alexander Lopez, M.S., John D. Overton, Ph.D., Thomas D. Schleicher, M.S., Maria Sotiropoulos Padilla, M.S., Karina Toledo, Louis Widom, Sarah E. Wolf, M.S., Manasi Pradhan, M.S., Kia Manoochehri, Ricardo H. Ulloa

**Genome Informatics**

Xiaodong Bai, Ph.D., Suganthi Balasubramanian, Ph.D., Leland Barnard, Ph.D., Andrew L. Blumenfeld, Yating Chai, Ph.D., Gisu Eom, Lukas Habegger, Ph.D., Young Hahn, Alicia Hawes, Shareef Khalid, Jeffrey G. Reid, Ph.D., Evan K. Maxwell, Ph.D., John Penn, M.S., Jeffrey C. Staples, Ph.D., Ashish Yadav, M.S.

**Clinical Informatics**

Nilanjana Banerjee, Ph.D., Michael Cantor, M.D.

**Analytical Genomics and Data Science**

Goncalo Abecasis, Ph.D., Lauren Gurski, Alexander Li, Ph.D., Daren Liu, Jonathan Marchini Ph.D., Anthony Marcketta, Shane McCarthy, Ph.D., Colm O'Dushlaine, Ph.D., Claudia Schurmann, Ph.D., Dylan Sun, Tanya Teslovich, Ph.D., Cristopher Van Hout, Ph.D., Bin Ye

**Therapeutic Area Genetics and Pharmacogenomics**

Giovanni Cappola, M.D., Amy Damask, Ph.D., Jan Freudenberg, M.D., Nehal Gosalia, Ph.D., Claudia Gonzaga-Jauregui, Ph.D., Julie Horowitz, Ph.D., Nan Lin, Ph.D., Charles Paulding, Ph.D., Kavita Praveen, Ph.D.

**Functional Modeling**

Shek Man Chim, Ph.D., Giusy Della Gatta, Ph.D., Valerio Donato, M.D./Ph.D., Aris Economides, Ph.D., Wan-Ying Hsieh, Ph.D., Lawrence Miloscio, Harikiran Nistala, Ph.D., Trikaldarshi Persaud, Chris Schoenherr, Ph.D.

**Planning, Strategy, and Operations**

Paloma M. Guzzardo, Ph.D., Marcus B. Jones, Ph.D., Lyndon J. Mitnaul, Ph.D.
